# Protein search processes mediated by chromatin topology

**DOI:** 10.1101/2023.12.12.571394

**Authors:** Shuvadip Dutta, R. Adarshkrishnan, Ranjith Padinhateeri, Mithun K. Mitra

## Abstract

We investigate the role of compaction of chromatin domains in modulating search kinetics of proteins. Collapsed conformations of chromatin, characterised by long loops which bring distant regions of the genome into contact, and manifested structurally as Topologically Associated Domains (TADs) affect search kinetics of DNA associated transcription factors and other proteins. In this study, we investigate the role of the compactness of chromatin on the dynamics of proteins using a minimal model. Using analytical theory and simulations, we show that an optimal compaction exists for which the residence time of proteins on a chromatin-like polymer backbone is minimum. We show that while bulk diffusion is an advantageous search strategy for extended polymers, for highly folded polymer domains, intersegmental transfers allow optimal search. We extend these results to more detailed polymer models - using the Freely Rotating Chain model, a Lennard-Jones bead-spring polymer model, which approximates chromatin behavior. We show that our results continue to hold for these polymer models, with a minimum residence time at an optimum polymer compaction. Finally, we also analyse the dynamics of proteins on networks generated using experimental chromatin conformation data from 8355 TADs extracted from human chromosomes. Our analysis suggests that TADs exist near this zone of optimality, indicating that chromatin conformations can play a crucial role in modulating protein search strategies.

## I. INTRODUCTION

The mechanism of target search by DNA-associated proteins such as Transcription Factors (TFs) on DNA backbones remains an open question [1–4]. Simple three-dimensional diffusion to locate specific binding sites cannot account for search timescales, necessitating proposals of facilitated diffusion processes [5–8]. Four distinct search mechanisms have been proposed to explain the dynamic motion of proteins along DNA: 1D sliding, hopping or jumping, 3D diffusion, and intersegmental transfer [9]. A combination of 3D and 1D diffusion has been shown to lead to optimal search times compared to a pure 1D or a pure 3D search [5]. However, experimental evidence points to a preponderance of sliding modes of transport, with only very small time spent in the bulk (SI Table S1) [10, 11]. In contrast, intersegmental hops or “direct transfers” have been observed experimentally [12], especially in densely packed nucleoids. This assumes particular significance for protein motion in mammalian cells, where highly packed chromatin regions with high density and viscosity hinder free diffusion in the solution phase [13], and multivalent proteins, due to nonspecific interactions, slow the mobility of other searching proteins, limiting their capacity for free 3D diffusion [14].

In this crowded milieu, intersegmental transfer from one segment of DNA to another via long loops assumes particular significance as a mechanism for effective search, which does not include free diffusion in the bulk [9]. While most studies assume that intersegmental transfer moves the protein to an uncorrelated segment, and thus has the same effect as 3D diffusion [2], this is not true for densely packed chromatin, where backbone correlations persists over long length scales, and impact the intersegmental transfer dynamics. Numerous theoretical and experimental studies have delved into the hierarchical organization of the genome across various length scales [15–21]. In mammals, the genome is organised into compact regions, such as Topologically Associating Domains (TADs) [22–24]. TADs are structural and functional units of the genome characterised by large squares of increased contact frequency tiling the contact matrices’ diagonal [23] (Fig. 1a). Concurrent with this high intra-domain contacts inside TADs, is a relative insulation among adjacent domains through TAD boundaries [25, 26]. TAD boundaries, marked by the enrichment of H3K4Me3 and the presence of abundant Pol2 [27], serve as hubs where the promoter regions of numerous transcriptional units are strategically located. These enhanced contacts within TADs result in a higher occurrence of long intersegmental hops, consequently influencing t he kinetics of the search processes of proteins on chromatin [28]. Indeed, search processes on polymer backbones have shown surprising results for protein kinetics [8], both in presence and absence of polymer fluctuations [29, 30]. DNA conformations and looped structures such as rosettes have been predicted to affect search kinetics [31–33].

**FIG. 1.**
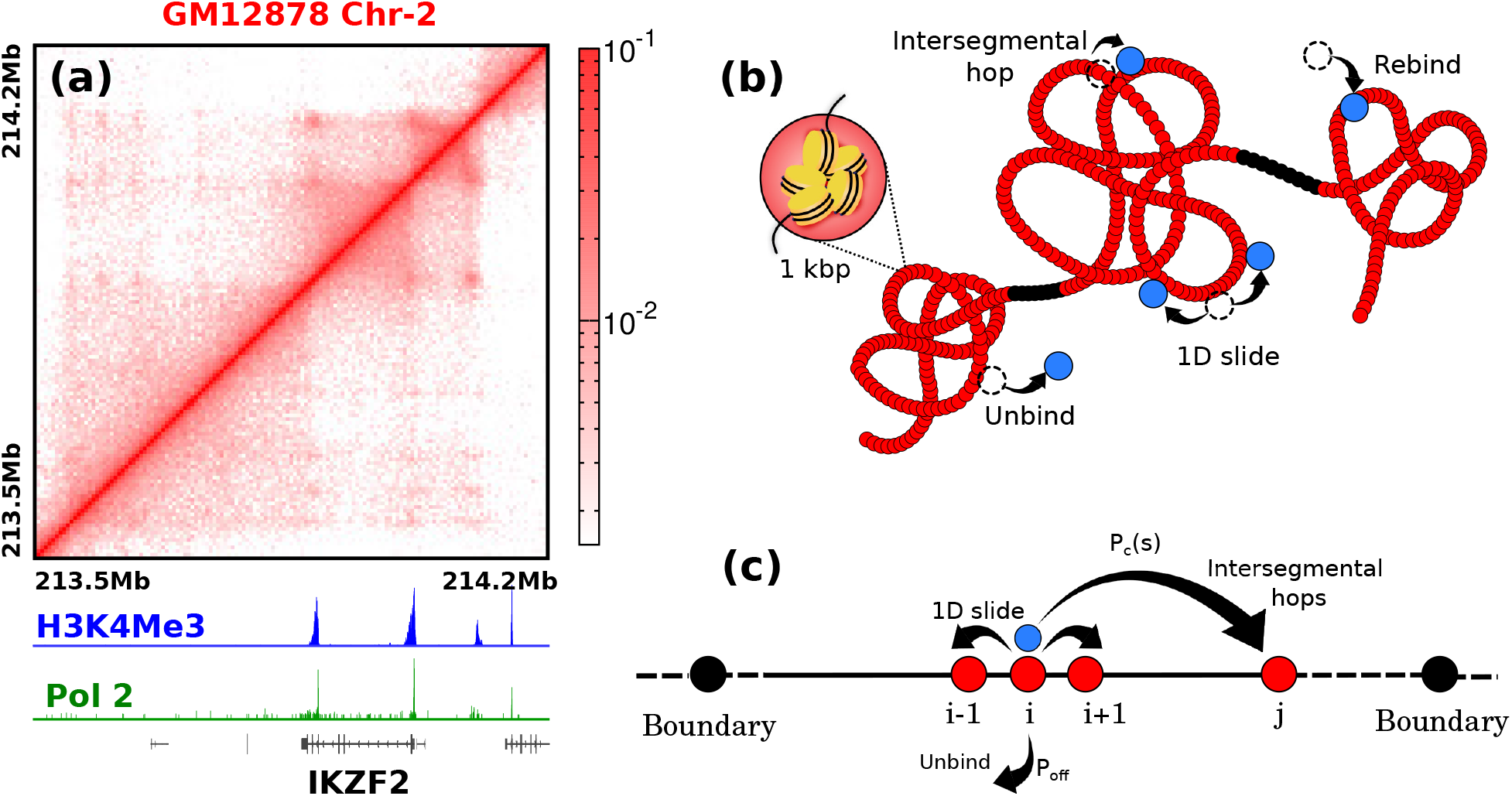
**(a)** A representative contact domain shows TADs as square blocks of increased interaction frequencies [34]. The appearance of a TAD is often accompanied by H3K4Me3 and Pol-2 enrichment peaks at the boundaries, as well as the activation of a gene whose promoter lies at one of the peak loci. **(b)** Coarse-grained (1 kbp) polymer representation of chromatin with compact TAD regions (red) separated by relatively extended boundary regions (black). **(c)** A contact space representation of TAD-like polymer domains. Each bead is connected via bonds to its neighbor along the backbone. In addition, bonds between genomically distant but spatially neighbouring beads allow for intersegmental jumps. Black beads denote the boundaries of the TAD.

In this Letter, we investigate the role of 3D chromatin organization, in particular within folded TAD-like regions, in the target search process by proteins. We show how the search times of proteins crucially depend on the topology of polymer domains characterised by long-range intersegmental contacts. In biologically relevant regimes, we show that 3D diffusion is not a efficient search strategy compared to search mediated through intersegmental jumps. Finally, we analyse available experimental data to show that conformations of human TADs are optimised for efficient search.

## II. A NETWORK REPRESENTATION OF CHROMATIN

We first consider a contact space network representation of a polymer domain (SI Sec. 1.1, Fig. S1). Two beads *i* and *j* (1 ≤ *i, j* ≤ *L* − 1) are assumed to be in contact with a probability *P*_*c*_(*s*) = *cs*^−*γ*^, where *s* = |*i* − *j*| *>* 1. A protein performs a random walk (SI Sec. 1.2) along the contact space, and can also dissociate to the bulk from any site with a probability *p*_off_ (Fig. 1 b,c). The target sites are located just outside of this polymer domain, at *i* = 0, *L*, motivated by the presence of transcriptional hubs at the TAD boundaries (Fig. 1a). A protein in the bulk can attach randomly to any site of the polymer, including the target sites, with a mean exploration time of *τ*_*f*_ in the solution phase before it reattaches to the polymer.

We compute the configuration-averaged mean search times ⟨*T*_search_⟩, defined as the time a protein takes to reach either boundaries of the polymer for the first time (SI Sec. 1.3).

The mean search time can be computed as

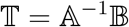

where 𝕋_*i*_ is the time taken to reach the target starting from the *i*^*th*^ site. The matrices 𝔸 and 𝔹 are defined by,

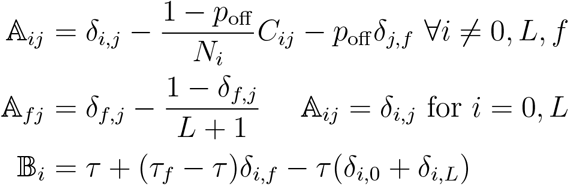

where, *τ* is the time taken for a single step, *C*_*ij*_ denotes the connectivity matrix, and *N*_*i*_ denotes the total number of bonds for the *i*^*th*^ site. Note that the polymer backbone connectivity implies that *C*_*i,i*−1_ = *C*_*i,i*+1_ = 1 necessarily. The site *i* = *f* denotes the freely diffusing state in solution. For simplicity, we assume the protein starts from the middle of the polymer domain, ⟨*T*_search_⟩ = ⟨𝕋_*L/*2_⟩.

Motivated by experimental results of very high estimates of time a protein spends non-specifically bound to the DNA [10, 11], and consistent with high volume fractions inside TADs [35] (SI Sec. 5,Fig. S12), we first consider the limiting case *p*_off_ = 0, so that target search happens only along the polymer backbone, via sliding and intersegmental jumps. For simplicity, we first assume *γ* = 0, so that all beads have a uniform connection probability *P*_*c*_(*s*) = *c* ≡ *p*_*u*_. We observe a non-monotonic relation between ⟨*T*_search_⟩ and *p*_*u*_ (Fig. 2a). When *p*_*u*_ = 0, the protein performs a 1D random walk, giving 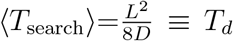 where 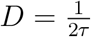 [36]. As *p*_*u*_ increases, more paths to reach the boundaries emerge, reducing ⟨*T*_search_⟩. However, beyond a threshold connectivity, ⟨*T*_search_⟩ increases due to excess intersegmental bonds diverting the walker. This non-monotonic behavior of ⟨*T*_search_⟩ remains robust even when the polymer network undergoes dynamic re-configurations (SI Sec. 2, Fig. S2) at a fixed connection probability (Fig. 2a red symbols, and SI Fig. S3a)). Dynamic rewiring of network connections reduces ⟨*T*_search_⟩, with faster configuration changes yielding lower ⟨*T*_search_⟩ at low *p*_*u*_ (SI Fig. S3a). However, at high *p*_*u*_ values, network dynamicity offers little advantage, as most nodes already have numerous long-range connections. The optimal behavior of ⟨*T*_search_⟩ is consistent for all polymer length (SI Fig. S4), differential rates for sliding and intersegmental jumps (SI Fig. S3b), and differential dynamic rewiring rates (SI Fig. S3a).

**FIG. 2.**
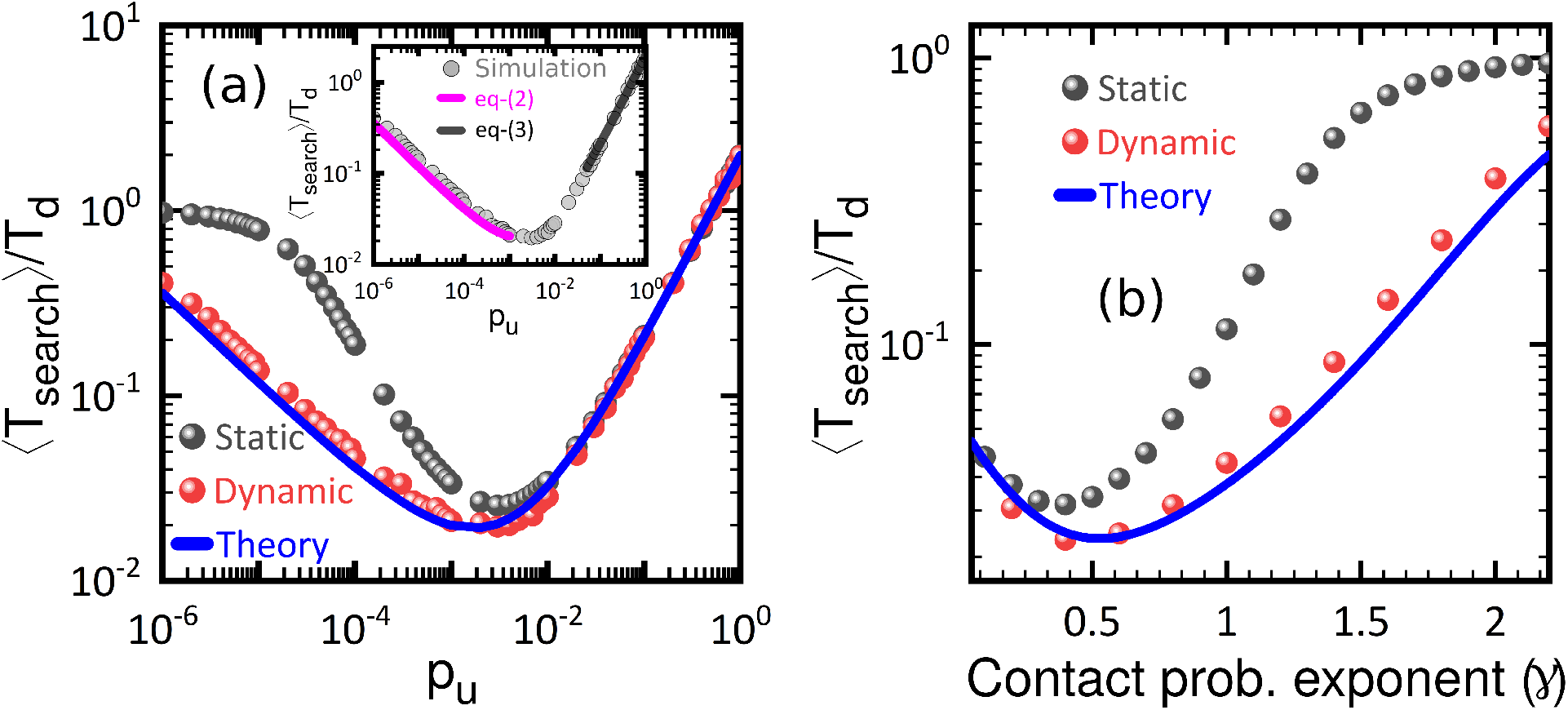
**(a)** Scaled mean search time 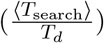 for static (black) and dynamic (red) conformations in a uniformly connected network domain vs probability of contact (*p*_*u*_) between any two non-neighbouring beads for *L* = 500. The solid blue line is the analytical solution of Eq. 1. The inset shows the low-*p*_*u*_ and high-*p*_*u*_ approximations from Eqs. 3 and 4. **(b)** 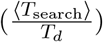 in power-law connected network domain as a function of contact probability exponent(*γ*) for *L* = 500 also shows non-monotonic behaviour for both static (black) and dynamic (red) configurations.

The non-monotonic behaviour persists when we use a power law connectivity with *γ*≠ 0. For a self-avoiding polymer, *γ* = 2.2, which reduces to *γ* = 1.5 for a RW polymer. For chromosomes, on an average, *γ* ≈ 1.08 [37], while within highly folded TADs, experiments report even smaller values for *γ* [38]. As a polymer undergoes collapse, contacts become more frequent, reducing the contact probability exponent *γ*. We observe a non-monotonic behaviour of the search times for both and static and dynamic realisations of such power-law networks, as shown in Fig. 2b. Again, when the polymer is relatively open (larger *γ*), dynamic rewiring of the connectivities helps reduce search times, similar to the observation from the simpler uniform connectivity (*P*_*c*_(*s*) = *p*_*u*_) case.

For the case of the dynamic polymer, where the polymer undergoes rewiring at a fixed connection probability *p*_*u*_, we can solve for the mean search times analytically. The mean search times (*T*_*k*_(*p*_*u*_)) for a protein starting from the *k*^th^ site obeys,

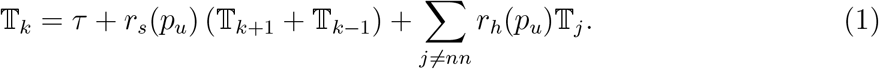

The total number of bonds for any site *i* is *N*_*i*_ = 2 +*p*_*u*_(*L*−3), and all hops are equally likely. The probability for sliding to any of the two neighbouring sites is then *r*_*s*_(*p*_*u*_) = 1*/N*_*i*_ and the probability of hopping to a non-neighbouring site via a intersegmental hop is *r*_*h*_(*p*_*u*_) = *p*_*u*_*/N*_*i*_ on average (SI Sec. 3). This can be solved exactly, subject to the absorbing boundary conditions at the two ends to yield a closed form solution (SI Sec. 3.2).

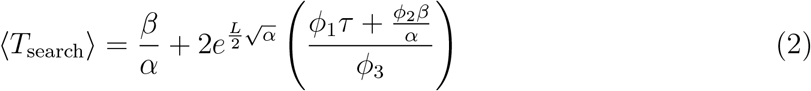

where

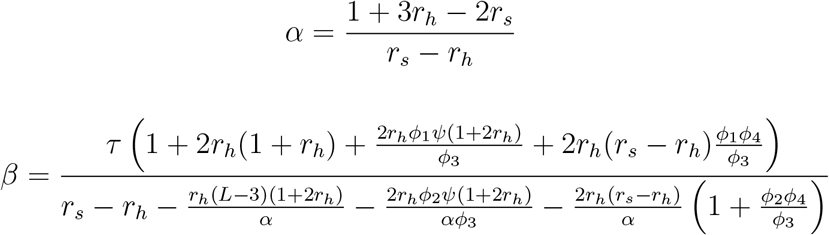

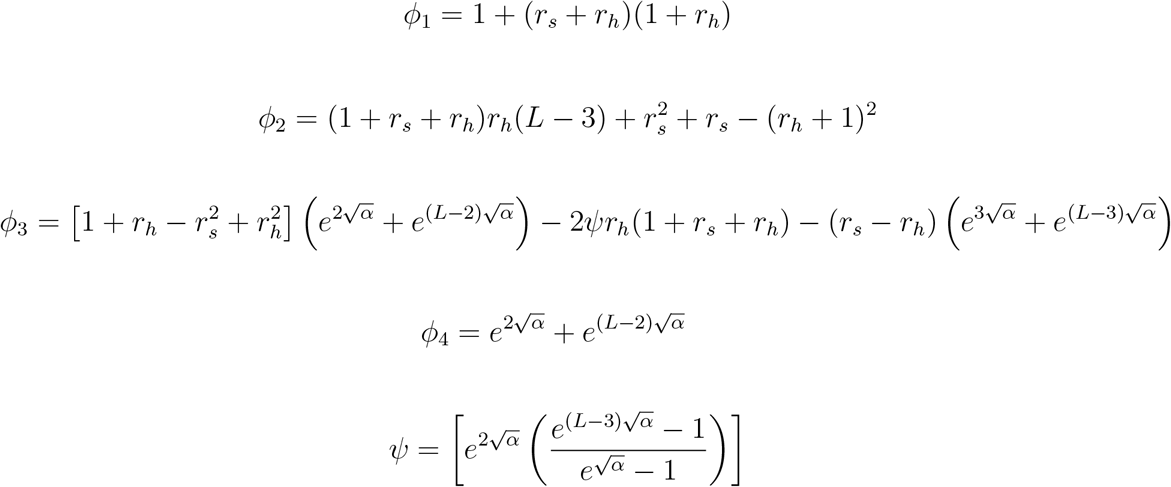

The analytic matches expression exactly with the simulation results (blue solid line in Fig 2a). For low values of *p*_*u*_, the governing equation can be simplified to obtain a solution (SI Sec. 3.2.1),

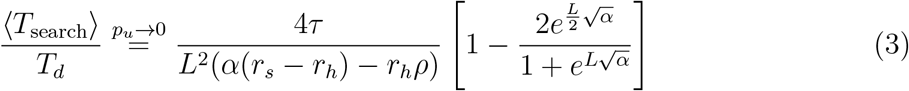

where, 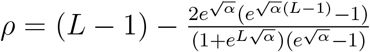 and 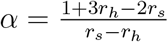 To leading order, then, we have,

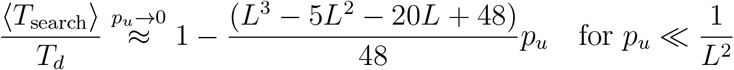

a decreasing function of *p*_*u*_. At high connection probability, as *p*_*u*_ → 1, all sites become equivalent, and a simplified governing equation can be solved to yield (SI Sec. 3.2.2),

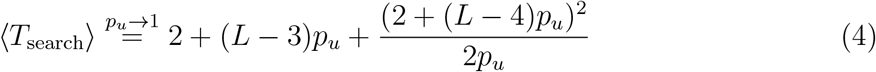

which implies a linear increase with *p*_*u*_ as *p*_*u*_ → 1, also consistent with a solution using the effective medium approach [39] (SI Sec. 3.2.3 and SI Fig. S5,S6). As can be seen from Fig 2a inset, the analytic expression at both small and high *p*_*u*_ limits matches the simulation results.

How does this search strategy mediated by intersegmental jumps compare with the facilitated diffusion in the presence of 3D diffusion (*p*_off_ ≠ 0)? We first look at the case of *γ* = 0 (Fig. 4a), so that *P*_*c*_(*s*) = *p*_*u*_. At *p*_*u*_ = 0, we recover the classical facilitated diffusion result of an optimum search at an intermediate value of *p*_off_. As *p*_*u*_ increases, the advantage obtained through bulk diffusion reduces, until beyond a certain threshold, unbinding from the backbone offers no advantage to the target search process. This general result holds true when we look at power-law connectivities (Fig. 4b) as well. For the self-avoiding polymer (*γ* = 2.2, 3D diffusion offers an advantage to target search. As the polymer collapses, and hence *γ* decreases, the relative speed-up through bulk excursions decreases. Beyond the RW polymer (*γ* ∼ 1.5), unbinding from the backbone offers no advantage to the target search process, as intersegmental hops allow a more efficient mechanism to explore distant sites. Indeed for collapsed polymers (low *γ* or large *p*_*u*_), and in the vicinity of the optimal compaction state, remarkably, 3D diffusion is not helpful for target search, with a monotonic increase in search times with increasing unbinding probability (SI Sec. 3.3, Fig. S7).

**FIG. 3.**
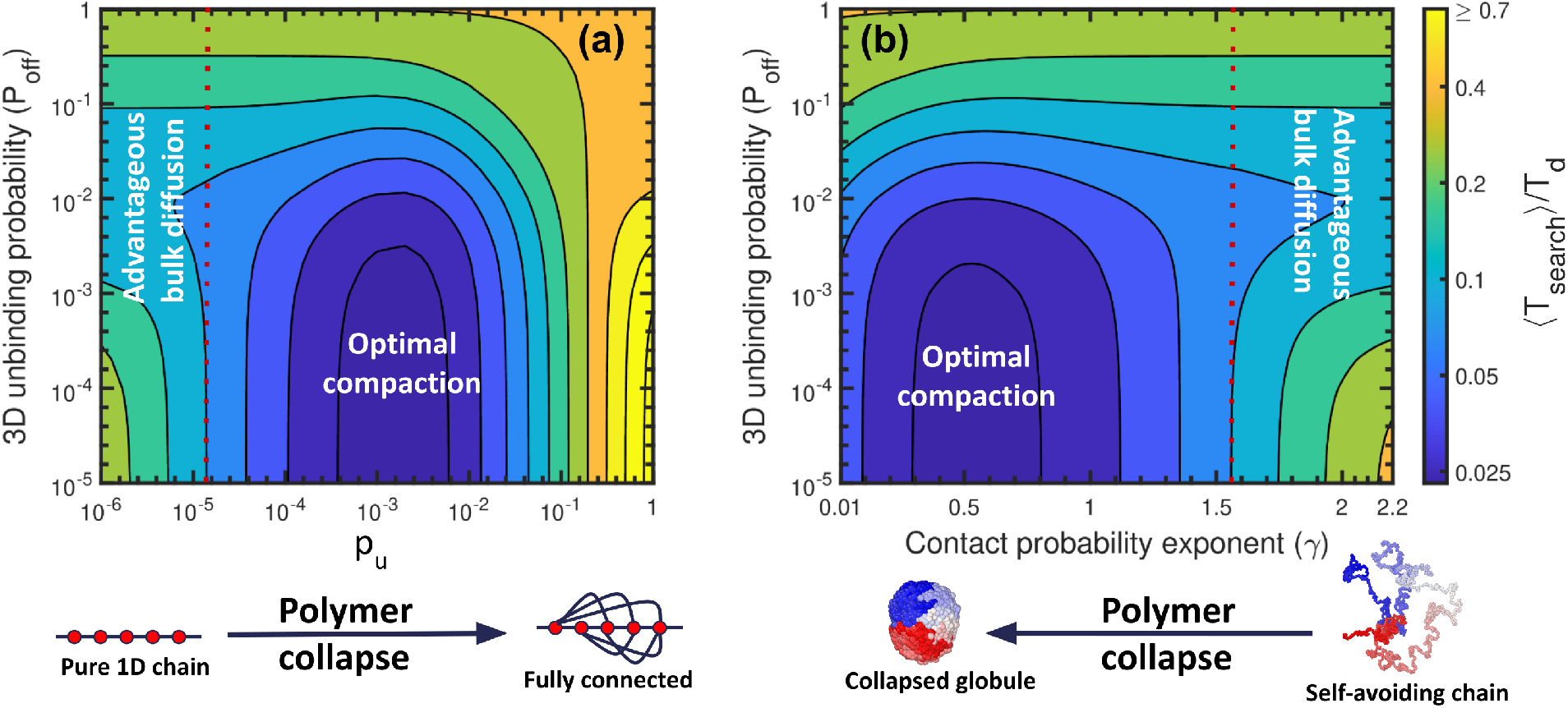
Contour plot of ⟨*T*_search_⟩*/T*_*d*_ as a function of 3D unbinding probability (*p*_off_) and polymer compaction for **(a)** a uniformly connected domain and **(b)** a power-law connected domain of *L* = 500. In a relatively open chain (see left side in **(a)** and right side in **(b)**, there is an optimal *p*_off_ for which ⟨*T*_search_⟩ is minimal suggesting advantage of 3D excursion. However, as polymer collapses, ⟨*T*_search_⟩ is minimum for the smallest *p*_off_ suggesting that 3D excursions are not advantageous. For biological *p*_off_ regimes, there always exists an optimal polymer compaction where ⟨*T*_search_⟩ is minimum. The fact that the global minimum of ⟨*T*_search_⟩ occurs for highly folded conformations and in regions where unbinding is low suggests the role of 3D structure in search process (see text).

**FIG. 4.**
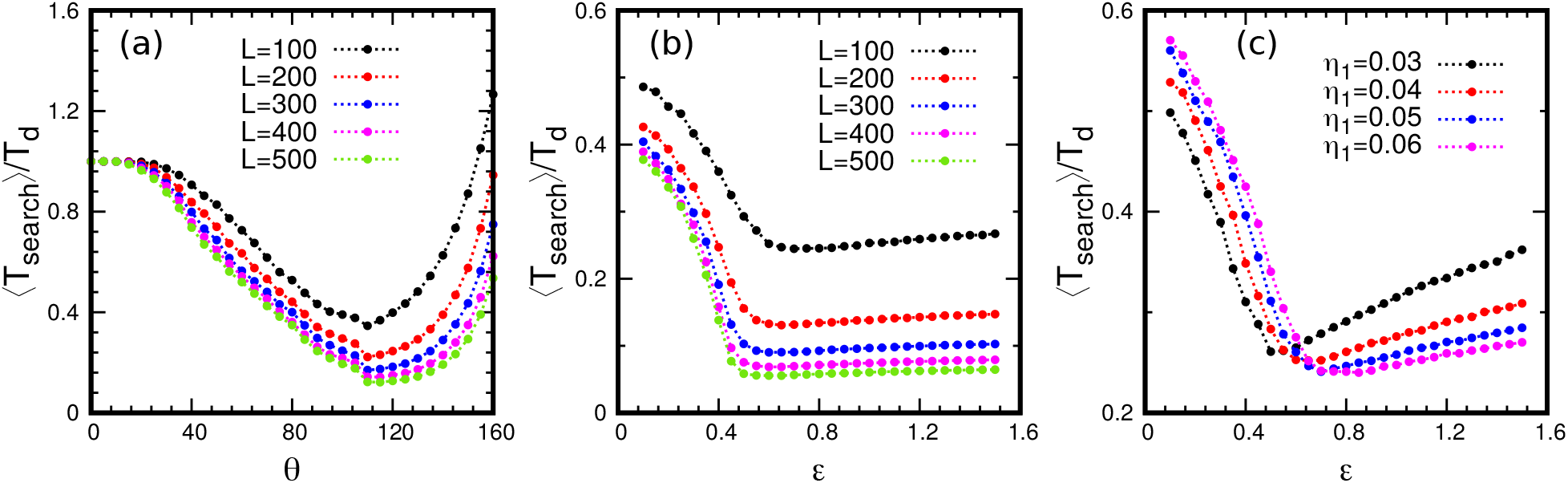
**(a)** Scaled mean residence time 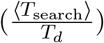 in FRC polymer domain as a function of angle (*θ*) between any two bond vectors for *L* = 100 − 500. **(b)** 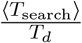 in LJ polymer domain as a function of LJ interaction strength (*ϵ*) between any two non-bonded beads for *L* = 100 − 500. **(c)** 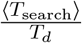 in soft LJ polymer domain as a function of soft LJ interaction strength (*ϵ*) between any two non-bonded beads for *L* = 100 with varying softness characterised by *η*_1_.

## III. POLYMER REPRESENTATIONS OF CHROMATIN

We now examine whether these effects persist when canonical polymer models are used to model chromatin structure. Thus the allowed intersegmental jumps arise directly from polymer topology, and hence random walks on these backbones have direct relevance to protein search processes.

### A. Freely Rotating Chain Model

We first study the Freely Rotating Chain (FRC) model which has been used to investigate chromatin behaviour [40–43]. Neighbouring bonds in an FRC polymer are constrained to lie on a cone subtending an angle *θ* [44]. The angle *θ* controls the compaction of the polymer, with large *θ* leading to a collapsed state (see SI Fig. S8A). As the polymer collapses, the number of non-neighbouring bonds *N*_*b*_ increases monotonically (see SI Fig. S8B), indicating a greater probability of intersegmental jumps. We simulated search on these FRC backbones (see SI Sec. 4.1), and the mean residence times ⟨*T*_search_⟩ shows a non-monotonic behaviour with increasing bond angle (Fig. 4A). This can be also interpreted as a function of polymer compactness or collapse. Thus in line with the expectations from the network model of polymer, there exists an optimal compaction where residence times are minimum.

### B. Lennard-Jones polymer model

We next use a bead-spring polymer model of chromatin with attractive Lennard-Jones (LJ) potential [45–47]. This introduces self-avoidance effects, and bond length fluctuations absent in the FRC model. Any pair of beads of diameter *σ*, separated by distance *r*, interacts via the potential,

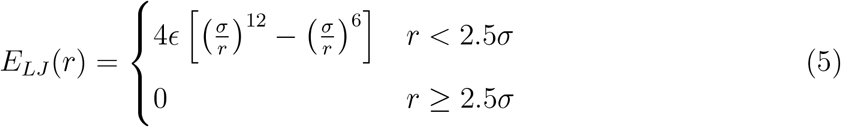

where *ϵ* is the LJ interaction strength (see SI Sec. 4.2). A LJ polymer undergoes a second-order phase transition from a coiled to a compact state (See SI Fig. S9A), accompanied by an increase in the number of intersegmental bonds (see SI Fig. S9B), as *ϵ* is increased. The residence times ⟨*T*_search_⟩ on these LJ polymer backbones exhibited a non-monotonic pattern as the LJ interaction strength is increased (Fig. 4B), or equivalently as the polymer compaction increases. There was a distinct minimum observed in the residence times, occurring close to the collapse transition. Thus a LJ polymer qualitatively mirrors the results obtained from the uniform network and FRC polymer models.

However, unlike the significant rise seen in the FRC and the network model, the extent of ⟨*T*_search_⟩ increase beyond a critical *ϵ* is less pronounced in this case. This limited increase is due to the inherent hard-core repulsion in the LJ potential, which imposes a lower limit on polymer compaction, and on the number of intersegmental bonds. This interplay between LJ interactions and polymer structure thus introduces a unique feature that distinguishes it from both FRC and the network model.

### C. Soft Lennard-Jones polymer model

Is this weak non-monotonic rise in ⟨*T*_search_⟩ then the correct physical description of residence times? The answer hinges on whether hardcore LJ potential best represents coarse-grained chromatin conformations. Experimental and theoretical studies of chromatin configurations and 3D distances suggest that a soft inter-bead potential that allows for overlap is a more realistic description of coarse-grained chromatin [20, 48]. We study a polymer model with following soft inter-bead potential,

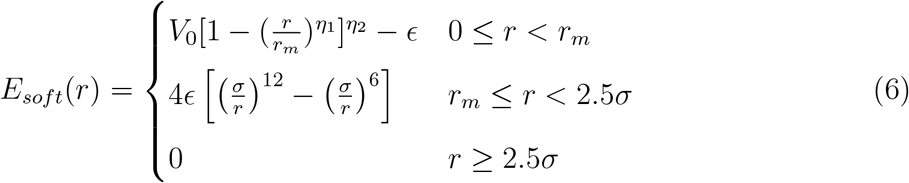

where *V*_0_ − *ϵ* is the energy penalty at complete overlap, and *η*_1_ and *η*_2_ tune the softness of the potential (see SI Sec. 4.3 and SI Fig. S10). We investigate ⟨*T*_search_⟩ in polymers with varying degrees of softness by tuning *η*_1_. A lower value of *η*_1_ indicates a softer polymer, whereas high *η*_1_ resembles the classical LJ potential. For a very soft polymer (*η*_1_ = 0.03), ⟨*T*_search_⟩ shows a strong non-monotonicity, analogous to the FRC and the network results. This is because a soft polymer core cannot effectively enforce volume exclusion, resulting in a more tightly packed polymer compared to the conventional LJ model (see SI Fig. S11A). This then leads to a higher number of non-neighboring contacts within the polymer domain (see SI Fig. S11B). As we decrease the softness (increasing *η*_1_), the degree of non-monotonicity in the residence times decreases and approaches the LJ-like behaviour. Thus non-monotonic behaviour of residence times remains a crucial feature of protein diffusion on coarse-grained chromatin-like soft polymer topologies. The degree of non-monotonicity ⟨*T*_search_⟩ is controlled by the softness of the inter-bead potential.

## IV. POLYMERS FROM TAD DATA

Is this non-monotonic behaviour of search times relevant for real TADs? Where do TAD conformations lie with respect to this optimal compaction state? To answer these, we utilized publicly accessible experimental Hi-C contact probability data [34] and performed the following theoretical study. We analysed Hi-C data for 8355 TADs at 1kbp resolution from GM12878 cell line Fig. 5a. For each TAD of length *L*, the contact probability curve was fitted to the equation *P*_*c*_(*s*) = *cs*^−*γ*^ in the region *s* = 10 to (*L* − 100) kbp Fig. 5b. The distributions of the best fitting values of *c* and *γ* are shown in 5c,d and yield ⟨*c*⟩ = 0.0214 and ⟨*γ*⟩ = 0.71 (most probable gamma values are in the range ≈ (0.66, 0.75)) (SI Sec. 6, Fig. S13). We computed protein target search times for wide range of *L* and *γ* on a power-law network with *c* = ⟨*c*⟩ and is shown as contour map in 5e. Strikingly, when we plot the best fitting *γ* values along with the length *L* for each TAD in the same Fig. 5e we find that they fall near the optimal search times (see red dots). This suggests that TAD structures for all human TADs are optimised for efficient search.

**FIG. 5.**
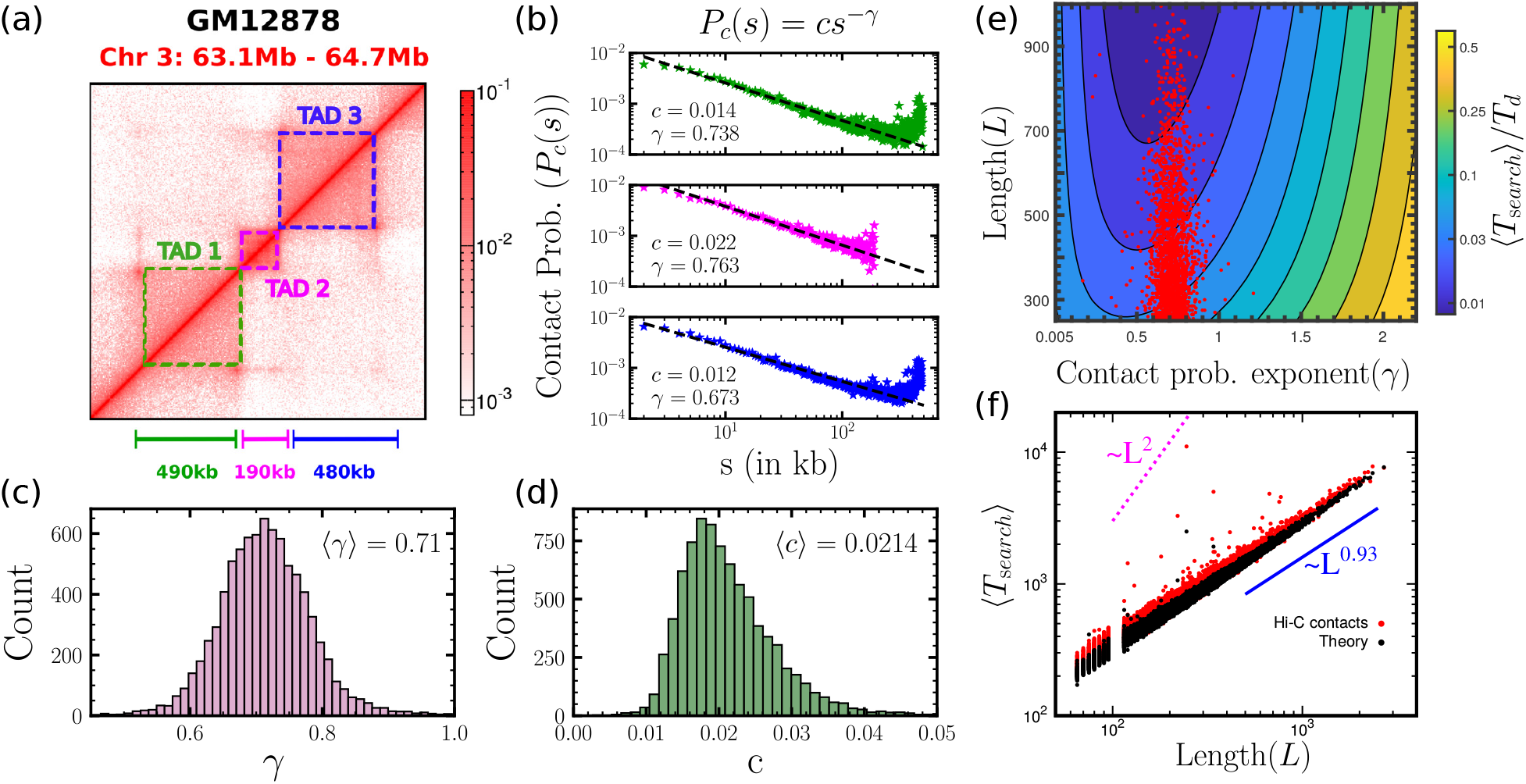
**(a)** An example of 3 distinct TAD regions in Hi-C contact map from [34]. **(b)** Contact probability (*P*_*c*_(*s*)) as a function of genomic distance (*s*), fitted using *P*_*c*_(*s*) = *cs*^−*γ*^ for the three TADs in (a). **(c-d)** Distribution of best fit parameters *c* and *γ* for all 8355 collected TADs from GM12878 data [34]. **(e)** Contour plot of mean search times (scaled) as a function of *γ* and *L*. The fact that experimentally observed [34] *L* and *γ* values (red dots) fall very close to the minimum is yet another evidence that 3D chromatin structure is relevant in the search processes. **(f)** Search times with TAD lengths show a very different scaling compared to purely 1D diffusive search. Search time computed from simulating walks on network with Hi-C contact probability (red dots) and network with observed power law properties (black) show ⟨*T*_search_⟩ ∼ *L*^0.93^.

To investigate this further we generated chromatin network by following two different procedures: (i) power-law networks for each TAD of length *L* by taking the best fitting *c* and *γ* values from above, and (ii) direct Hi-C networks for all TADs by inserting connections directly with experimentally measured contact probabilities *P*_*ij*_. While the former gives accurate average contact probability for each *s* = |*i*−*j*|, the latter gives us accurate sequence-specific contact probability for every pair (*i, j*). We performed the search simulations on both networks and the results are shown in Fig. 5f. Note that, both the results (red,black) give comparable search times and the scaling with TAD lengths is close to ballistic (⟨*T*_search_⟩ ∼ *L*^0.93^). This implies that the TADs represented by red dots in Fig. 5e not only have the appropriate (*L, γ*) but also have the optimal search times. This suggests that random walks on these TADs can lead to large speed-ups compared to the diffusive timescales (∼ *L*^2^) purely due to the underlying TAD structure (Fig. 5f). Further, despite sequence-specific effects, search times on these Hi-C networks is accurately captured by power-law network models, and they offer a true estimate of times expected for search processes on chromatin backbones.

## V. DISCUSSION

In conclusion, we propose that facilitated diffusion which takes into account correlated intersegmental jumps can leverage the compaction state of chromatin, characterized by the presence of TADs and looped polymer conformations, to access distant genomic regions for more efficient search. This is in line with direct evidence from single-molecule experiments which have reported that search process by a site-specific restriction enzyme on DNA becomes almost doubly efficient when the DNA configuration is collapsed instead of a fully extended configuration [49]. Remarkably, contrary to the prevailing picture of bulk diffusion mediated search, we show that for highly folded polymer conformations, such as in chromatin, bulk diffusion hinders the search process, possibly offering a reconciliation with experimental estimated of very large times a protein spends non-specifically bound to DNA. An analysis of connection probabilities from experimentally determined Hi-C data suggests that TADs reside near this zone of optimal compaction where search times are minimised. The structural organization of the genome within TADs may thus have evolved to facilitate rapid and precise dynamics of DNA-associated proteins, which is crucial for gene regulation and cellular function.

## Supporting information

SI-Protein_search_processes_mediated_by_chromatin_topology.pdf

## VI. ACKNOWLEDGEMENT

SD acknowledges fellowship support from the PMRF, MoE, India. MKM and RP acknowledge funding from Core Research Grant by Science and Engineering Research Board, India (Grant number: CRG/2022/008142).

## Notes

### Competing Interest Statement

The authors have declared no competing interest.

### Summary of Updates

All figures revised; New results added in section 2; Detailed new analysis of search processes on Human TADs in section 4; Supplemental files updated.

