## Supplementary material for "Protein search processes mediated by chromatin topology": SI-Protein_search_processes_mediated_by_chromatin_topology.pdf

#### Contents

**S1 Methods** **S2**
S1.1 Contact space representation of protein search in chro- matin domain . . . S2
S1.2 Protein as random walker . . . S2
S1.3 Simulation details . . . S3

**S2 Robustness of the optimal compaction results** **S4**

**S3 Theoretical estimation of mean search times** **S5**
S3.1 Matrix inversion to get mean search times . . . S5
S3.2 Closed form solution for $p_{\text{off}} = 0$ . . . S5
S3.2.1 Small $p_u$ approximation . . . S10
S3.2.2 Large $p_u$ approximation . . . S11
S3.2.3 The effective medium approach at large $p_u$ . . . S12
S3.3 The effect of 3D unbinding on mean search time . . . S14

**S4 Search process on FRC, LJ and soft LJ polymer** **S15**
S4.1 Freely rotating chain model . . . S16
S4.2 LJ bead-spring polymer model . . . S16
S4.3 Soft LJ bead-spring polymer model . . . S17

**S5 Protein motion in coarse-grained chromatin** **S19**

**S6 Dynamics of protein in network constructed using Hi-C data** **S21**

### S1 Methods

#### S1.1 Contact space representation of protein search in chro- matin domain

We propose a theoretical framework for modeling a compact polymer domain of length $L$ as a network in contact space. In this model, each bead (node) within the polymer represents a coarse-grained chromatin segment of size 1 kbp. The conformation of the polymer is encoded using a matrix $c_{ij}$ of dimensions $(L+1) \times (L+1)$, where $i, j \in [0, L]$. In the network representation, the polymer backbone connectivity is considered, and additional connections between non-neighbouring beads are introduced. The existence of a connection between beads $i$ and $j$ for a given configuration is defined by $c_{ij}$ which takes the value 1 if a connection exists and 0 otherwise, the value taken by $c_{ij}$ is determined by a connection probability $P_c(s) = cs^{-\gamma} \in [0, 1]$ for all $i, j \in [1, L-1]$. It is worth noting that the polymer topology enforces $c_{i,i+1} = 1$ and $c_{i-1,i} = 1$ for all $i$. The exponent $\gamma$ serves as a measure for polymer compaction. For a self-avoiding polymer, $\gamma = 2.2$, which reduces to $\gamma = 1.5$ for a RW polymer. For chromosomes, on an average, $\gamma \approx 1.08$ [12], while within highly folded TADs, experiments report even smaller values for $\gamma$ [5]. As a polymer undergoes collapse, contacts become more frequent, reducing the contact probability exponent $\gamma$.

#### S1.2 Protein as random walker

In this network representation, a protein is depicted as an unbiased random walker navigating within a polymer domain $(i, j \in [1, L-1]$. The targets are located just outside the domain at $i = 0, L$. During each step (time $\tau$), a protein situated at bead $i$ can either slide along the chain to one of its 1D neighbors ($i+1$ and $i-1$) or hop to a genomically distant bead $j$ (3D neighbors) if $c_{ij} = 1$. Additionally, the walker can unbind with a probability $p_{\text{off}}$ to the bulk and subsequently rebind to the polymer randomly at any site, including its boundaries, after spending time $\tau_f$ in the bulk/solution phase. The boundaries of the polymer domain $i = 0$ or $i = L$ are absorbing. Upon reaching either of these boundaries, the walker is considered to have reached the target sites.

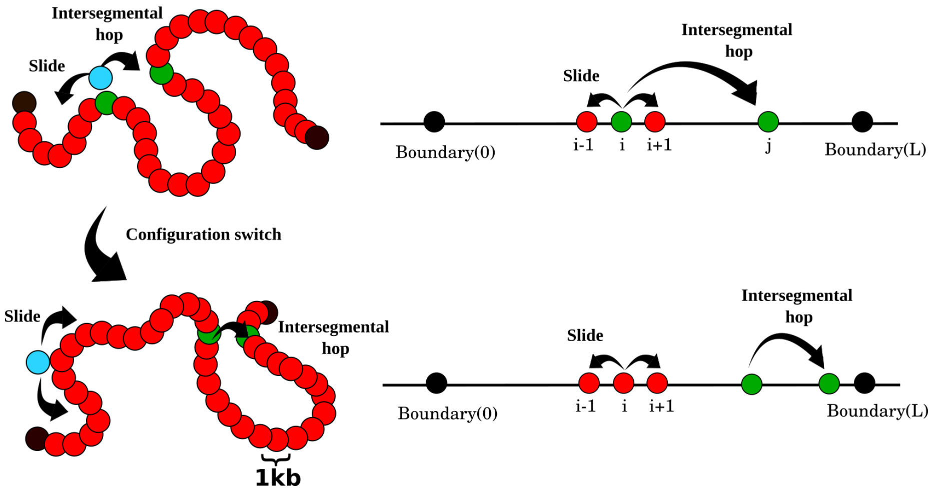


Figure S1: The polymer configuration (left) is depicted as a contact space network (right) where adjacent beads are connected to reflect the polymer connectivity. Additionally, non-adjacent beads have connections to represent 3D proximity within the polymer configuration. A protein, modeled as a random walker, can slide to either of its 1D neighbors along the chain or make jumps to available 3D neighbors, all with equal probability. The dynamic nature of the polymer allows for alterations in the polymer configuration, leading to the rewiring of network structures.

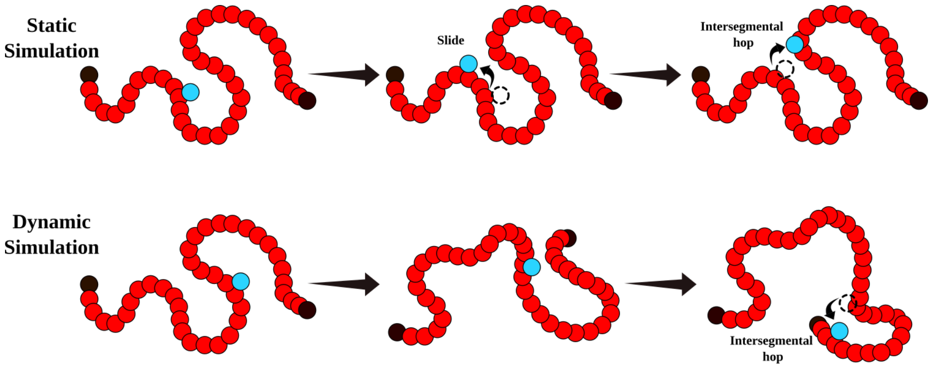


Figure S2: **Static and dynamic simulation:** In the static simulation (top), the network structure representing the polymer configuration remains constant throughout a single realization. The set of long-range connections is maintained without alteration. In contrast, in a dynamic simulation (bottom), the network structure undergoes continuous reconfiguration at a constant rate. This dynamic simulation is designed to capture the changing nature of the polymer configuration over time.

#### S1.3 Simulation details

We consider two cases depending on whether the connectivity matrix is allowed to evolve during the target search process. If the $c_{ij}$ do not change this represents the target search is happening on a static polymer configuration. On the other hand, if $c_{ij}$ evolve with time, this represents target search happening on a dynamic polymer. We describe the methodology of simulation for these two cases below.

##### Static domain

Given a contact probability $P_c(s)$ we construct a connectivity matrix $c_{ij}$. To determine the mean search time ($\langle T_{\text{search}} \rangle$) within the constructed domain, the random walker (protein) is initially positioned at the midpoint of the domain ($i = \frac{L}{2}$). The time taken for the walker to reach any of the absorbing boundaries ($i = 0, L$) for the first time is recorded. The mean search time ($\langle T_{\text{search}} \rangle$) is then computed by averaging the recorded times over ensembles (different realisations of $c_{ij}$ and trajectories.

#### Dynamic domain

In our previous analysis, we assumed a static domain, meaning that the configuration remained constant over time through out the search process. However, it is now widely recognized that chromatin organization within a cell is not a static structure but rather a dynamic polymer [8]. This dynamic nature arises from various factors, including the tethering of chromatin to the nuclear lamina, interactions with different proteins, and the dynamic process of loop extrusion mediated by cohesin [13]. In order to explore the potential impact of dynamics on the mean search times, we maintain our domain representation as described earlier. However, we introduce polymer dynamics by periodically altering the configuration of the domain. This dynamic aspect involves shuffling the positions of non-neighboring bonds ($c_{ij} = 1$) at regular interval of time $t_{\text{config-switch}}$. When $t_{\text{config-switch}} = \tau$, we call the configuration is dynamic as in each step of the protein configuration of the domain alters. On the other hand, when $t_{\text{config-switch}} >> \tau$, we denote the configuration is static. In essence, this approach allows us to simulate the dynamic nature of chromatin and investigate whether these network dynamics have any effect on the mean search times.

#### S2 Robustness of the optimal compaction results

The non-monotonic behavior of $\langle T_{\rm search}\rangle$ remains robust even when the polymer network undergoes dynamic re-configurations at a fixed connection probability (Fig. S3a). Dynamic rewiring of network connections reduces $\langle T_{\rm search}\rangle$, with faster configuration changes yielding lower $\langle T_{\rm search}\rangle$ at low $p_u$ values. However, at high $p_u$ values, network dynamicity offers little advantage, as most nodes already have numerous long-range connections. In Fig. S3b, we show that the non-monotonic behavior of $\langle T_{\rm search}\rangle$ persists even when sliding along the backbone and jumps along intersegmental bonds have different rates. At low $p_u$, the $\langle T_{\rm search}\rangle$ remains independent of $t_{\rm jump}$, dominated by 1D sliding. Conversely, for higher $p_u$, $\langle T_{\rm search}\rangle$ decreases as the $t_{\rm jump}$ increases, favoring long hops over sliding. The optimal behavior of $\langle T_{\rm search}\rangle$ is consistent for all polymer lengths, as shown Fig. S4.

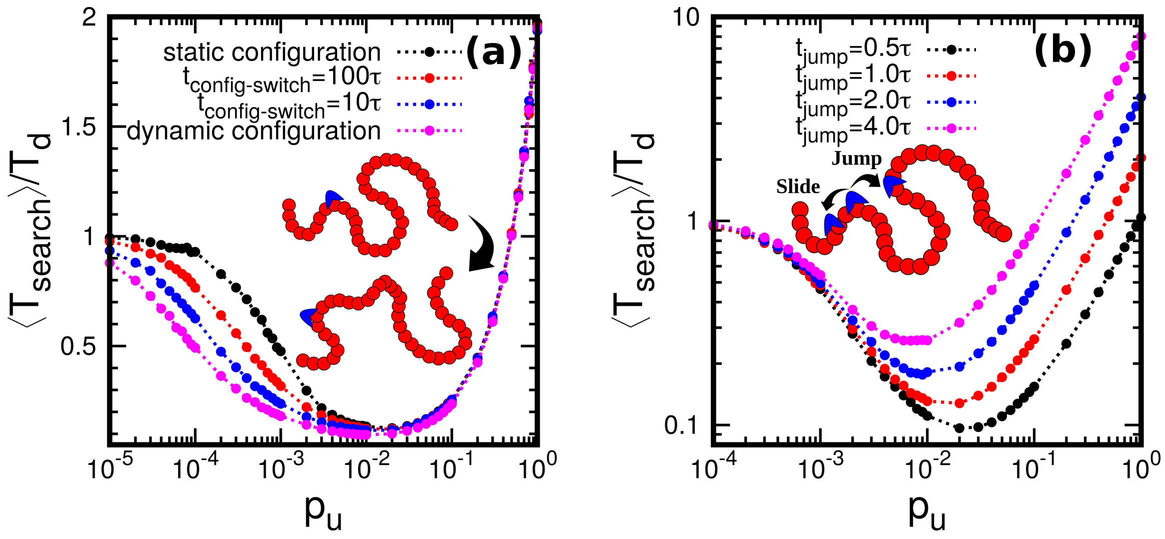


Figure S3: **(a)** $\frac{\langle T_{\rm search}\rangle}{T_d}$ in uniformly connected network domain vs $p_u$ for different dynamic rewiring for $L = 100$. **(b)** $\frac{\langle T_{\rm search}\rangle}{T_d}$ in uniformly connected network domain vs $p_u$ for different jump rates for $L = 100$.

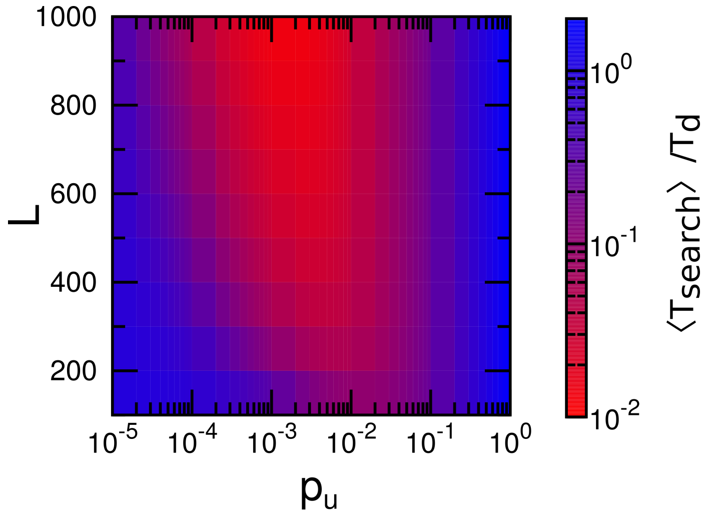


Figure S4: Mean search time in a uniformly connected domain as a function of $p_u$ and domain lengths ($L$). The colour bar represents scaled mean search time $\frac{\langle T_{\rm search}\rangle}{T_d}$.

#### S3 Theoretical estimation of mean search times

##### S3.1 Matrix inversion to get mean search times

The network model can be represented using a set of equations that can be solved using matrix inversion. We can formulate the target search process as a mean first passage time problem for the protein to reach the target sites at $0, L$ for the first time. The protein walking on the domain can unbind from any bead with a probability $p_{\text{off}}$ and rebind to any site including the boundary with a probability $\frac{1}{L+1}$ after spending a time $\tau_f$ diffusing in the bulk. At any site $i$, in addition to the nearest neighbours, protein can hop to 3D neighbours with a probability

$$P_c(s = |i-j|) = c|i-j|^{-\gamma} \quad \forall\, i, j \in [1, L-1]$$

When the walker is at position $i$ the dynamic average of possible connections

$$N_i = 2 + \sum_{j \neq nn, 0, L} c|i-j|^{-\gamma} \quad \forall\, i, j \in [1, L-1]$$

Assuming that the time to perform a slide/jump to the neighbouring bead is $\tau$, we can write the equation for average search time for a walker in the following matrix form

$$\mathbb{T} = \mathbb{A}^{-1}\mathbb{B}$$

where,

$$\mathbb{A}_{ij} = \begin{cases} \delta_{i,j} - \dfrac{1-p_{\text{off}}}{N_i}\big[\delta_{i\pm1,j} & \\ \quad + (1-\delta_{i\pm1,j}-\delta_{f,j})(1-\delta_{0,j}-\delta_{L,j})c|i-j|^{-\gamma}\big] - p_{\text{off}}\delta_{f,j}, & \text{if } i \neq f, 0, L \\ \delta_{i,j} - (1-\delta_{i,j})\dfrac{1}{L+1}, & \text{if } i = f \\ \delta_{i,j}, & \text{if } i = 0, L \end{cases}$$

$$\mathbb{B}_i = \tau + (\tau_f - \tau)\delta_{i,f} - \tau(\delta_{i,0} + \delta_{i,L})$$

##### S3.2 Closed form solution for $p_{\text{off}} = 0$

In the case where protein's motion is purely along the chromatin backbone i.e. unbinding probability $p_{\text{off}} = 0$, we can rewrite a master equation for $T_i$–

$$T_i = \tau + r_s T_{i+1} + r_s T_{i-1} + \sum_{j \neq i, i+1, i-1} r_h T_j \tag{S1}$$

Here $j$ is summed over all other possible non-neighbouring beads excluding boundaries$(0, L)$. $\tau$ is the time taken for a single step. As the walker takes no time to search if it starts from $0, L$, the absorbing boundary conditions for $T_i$ is given by-

$$T_0 = 0 \quad \text{and} \quad T_L = 0$$

In the bulk, $i \in (1, L-1)$, the number of allowed intersegmental jumps (excluding the two nearest neighbours and both the boundaries) is $L-4$. Thus, normalisation conditions for slide and jump probabilities are given by–

$$2r_s + \sum_{\substack{i'=1 \\ i' \neq i, i\pm1}}^{L-1} r_h = 2r_s + (L-4)r_h = 1 \quad \forall i \in (1, L-1)$$

Using these relations, we can calculate explicit values of $r_s$ and $r_h$ as a function of $p_u$ for a given $L$ in the domain $i \in (1, L-1)$,

$$r_s = \frac{1}{2 + p_u(L-4)} \qquad\qquad r_h = \frac{p_u}{2 + p_u(L-4)}$$

At $i = 1, L-1$, the number of allowed intersegmental jumps (excluding the two nearest neighbours and the opposite boundary) is $L-3$. Thus, the slide and jumps probabilities are denoted as $r'_s$ and $r'_h$ respectively and normalisation conditions are given by,

$$2r'_s + \sum_{\substack{i'=2 \\ i' \neq i \pm 1}}^{L-1} r'_h = 2r'_s + (L-3)r'_h = 1 \qquad\qquad \text{if} \quad i = 1$$

$$2r'_s + \sum_{\substack{i'=1 \\ i' \neq i \pm 1}}^{L-2} r'_h = 2r'_s + (L-3)r'_h = 1 \qquad\qquad \text{if} \quad i = L-1$$

which gives,

$$r'_s = \frac{1}{2 + p_u(L-3)} \qquad\qquad r'_h = \frac{p_u}{2 + p_u(L-3)}$$

We construct a differential equation for the problem in the domain $\Delta x < x < (L-1)\Delta x$, where $\Delta x$ is the spacing between two adjacent nodes and $L$ is the total number of nodes starting from zero. The near boundary deviance will be introduced into the closed-form solution using appropriately defined boundary conditions.

To make the summation term uniform for all $i$ values we add and subtract $r_h(T_{i-1} + T_i + T_{i+1})$ to the master equation(S1) to obtain-

$$(1 + r_h)T_i = \tau + (r_s - r_h)(T_{i+1} + T_{i-1}) + r_h \sum_{j=1}^{L-1} T_j$$

This can be rewritten as,

$$(1 + r_h)T_i - (r_s - r_h)(T_{i+1} + T_{i-1}) - r_h \sum_{j=1}^{L-1} T_j = \tau$$

The central difference formulation for the second derivative is given by-

$$\frac{\partial^2 T}{\partial x^2} = \frac{T_{i+1} - 2T_i + T_{i-1}}{(\Delta x)^2}$$

To introduce this form into our equation we add and subtract $2T_i$ to the second term hence giving us-

$$(1 + r_h)T_i - 2(r_s - r_h)T_i - (r_s - r_h)(T_{i+1} - 2T_i + T_{i-1}) - r_h \sum_{j=1}^{L-1} T_j = \tau$$

By multiplying and dividing the second term by $\Delta x^2$,

$$(1 + 3r_h - 2r_s)T_i - (r_s - r_h)\frac{(T_{i+1} - 2T_i + T_{i-1})}{\Delta x^2}(\Delta x^2) - (\tau + r_h \sum_{j=1}^{L-1} T_j) = 0$$

Assuming the given equation is in the finite difference form reverting it to the differential form while substituting $\Delta x = 1$ evaluates to-

$$(r_s - r_h)\frac{\partial^2 T[x]}{\partial x^2} - (1 + 3r_h - 2r_s)T[x] + (\tau + r_h \sum_{x=1}^{L-1} T[x]) = 0 \tag{S2}$$

We will concentrate on obtaining a closed-form solution for this equation within the bounds $0 < p_u < 1$ which says $(r_s - r_h) \neq 0$. We can rearrange the problem into the form-

$$\frac{\partial^2 T[x]}{\partial x^2} - \left(\frac{1 + 3r_h - 2r_s}{r_s - r_h}\right) T[x] + \left(\frac{\tau + r_h \sum_{x=1}^{L-1} T[x]}{r_s - r_h}\right) = 0 \tag{S3}$$

Since the summation term is independent of x we can prudently use the substitution:

$$\alpha = \frac{1 + 3r_h - 2r_s}{r_s - r_h} \qquad \beta = \frac{\tau + r_h \sum_{x=1}^{L-1} T[x]}{r_s - r_h}$$

to rewrite (S3) as-

$$\frac{\partial^2 T}{\partial x^2} - \alpha T + \beta = 0$$

The general solution of a differential equation of this form is given by-

$$T[x] = \frac{\beta}{\alpha} + C_1 e^{x\sqrt{\alpha}} + C_2 e^{-x\sqrt{\alpha}} \tag{S4}$$

Where $C_1$ and $C_2$ are constants of integration, we will use the boundary conditions to determine values of these constants.

**Condition 1:** $T[x]$ as a function of $x$ depending on the value of $p_u$ peaks at $x = \frac{L}{2}$ and going to zero at the boundaries. We can safely assume that the first derivative of the function $T[x]$ with respect to $x$ goes to zero at $\frac{L}{2}$. Taking the first derivative of $T[x]$ from (S11) and substituting the condition gives-

$$\Rightarrow \frac{\partial T[L/2]}{\partial x} = \sqrt{\alpha}\left(C_1 e^{\frac{\sqrt{\alpha}L}{2}} - C_2 e^{-\frac{\sqrt{\alpha}L}{2}}\right) = 0$$

from the above equation we can derive $\left[C_2 = C_1 e^{\sqrt{\alpha}L}\right]$ and substitute in (S11) to get-

$$T[x] = \frac{\beta}{\alpha} + C_1(e^{x\sqrt{\alpha}} + e^{(L-x)\sqrt{\alpha}}) \tag{S5}$$

**Condition 2:** We will use the information that $T[0] = 0$ and parabolic symmetry to find an alternate expression for $T[1]$ to substitute into the equation for $T[2]$ therefore allowing $T[2]$ to act as a proxy boundary. From the master equation, we have-

$$T[1] = \tau + r'_s(T[2]) + r'_h \sum_{x=3}^{L-1} T[x]$$

Where $r'_s$ and $r'_h$ are the values of $r_s$ and $r_h$ when $x \in \{1, L-1\}$. We will use the symmetry property to write $T[L-1] = T[1]$, we then add and subtract $r'_h(T[2])$ to transform the above equation into the form

$$(1 - r'_h)T[1] = \tau + (r'_s - r'_h)T[2] + r'_h \sum_{x=2}^{L-2} T[x]$$

The summation term can be evaluated as follows

$$\sum_{x=2}^{L-2} T[x] = \sum_{x=2}^{L-2} \left[\frac{\beta}{\alpha} + C_1(e^{x\sqrt{\alpha}} + e^{(L-x)\sqrt{\alpha}})\right]$$

$$= (L-3)\frac{\beta}{\alpha} + C_1(e^{2\sqrt{\alpha}} + e^{3\sqrt{\alpha}} + .... + e^{(L-3)\sqrt{\alpha}} + e^{(L-2)\sqrt{\alpha}} + e^{(L-2)\sqrt{\alpha}} + e^{(L-3)\sqrt{\alpha}} + .... + e^{3\sqrt{\alpha}} + e^{2\sqrt{\alpha}})$$

$$= (L-3)\frac{\beta}{\alpha} + 2C_1(e^{2\sqrt{\alpha}} + e^{3\sqrt{\alpha}} + .... + e^{(L-3)\sqrt{\alpha}} + e^{(L-2)\sqrt{\alpha}})$$

Using geometric progression series summation-

$$\sum_{x=2}^{L-2} T[x] = (L-3)\frac{\beta}{\alpha} + 2C_1 \left[e^{2\sqrt{\alpha}} \left(\frac{e^{(L-3)\sqrt{\alpha}} - 1}{e^{\sqrt{\alpha}} - 1}\right)\right]$$

We will denote-

$$\psi = \left[e^{2\sqrt{\alpha}} \left(\frac{e^{(L-3)\sqrt{\alpha}} - 1}{e^{\sqrt{\alpha}} - 1}\right)\right]$$

and rewrite the equation as-

$$\sum_{x=2}^{L-2} T[x] = (L-3)\frac{\beta}{\alpha} + 2C_1\psi \tag{S6}$$

Using (S6) the summation in $T[1]$ can be rewritten as

$$T[1] = \frac{1}{(1-r'_h)}\left[\tau + (r'_s - r'_h)T[2] + r'_h(L-3)\frac{\beta}{\alpha} + 2r'_hC_1\psi\right]$$

but $r'_h = \frac{p}{2+p(L-3)}$ and $r'_s = \frac{1}{2+p(L-3)}$, therefore

$$\frac{1}{1-r'_h} = \frac{1}{1-\frac{p_u}{2+p_u(L-3)}} = \frac{2+p_u(L-3)}{2+p_u(L-4)} = \frac{2+p_u(L-4)+p_u}{2+p_u(L-4)} = 1 + r_h$$

similarly we can derive

$$\frac{r'_s - r'_h}{1-r'_h} = r_s - r_h, \qquad \frac{r'_h}{1-r'_h} = r_h$$

Hence we can rephrase $T[1]$ as

$$T[1] = (1+r_h)\tau + (r_s - r_h)T[2] + r_h(L-3)\frac{\beta}{\alpha} + 2r_hC_1\psi \tag{S7}$$

We will be using the master equation of $T[2]$ to solve for $C_1$

$$T[2] = \tau + r_sT[1] + r_sT[3] + s\left(T[4] + T[5] + .... + T[L-2]\right) + r_hT[L-1]$$

To this equation we will add and subtract $r_h(T[2]+T[3])$ to obtain a summation term that we have already evaluated, we then use the property that $T[1] = T[L-1]$ to get

$$(1+s)T[2] = \tau + (r_s + r_h)T[1] + (r_s - r_h)T[3] + r_h\sum_{x=2}^{L-2} T[x] \quad \text{(make a substitution using (S6))}$$

$$= \tau + (r_s + r_h)T[1] + (r_s - r_h)T[3] + r_h(L-3)\frac{\beta}{\alpha} + 2r_hC_1\psi$$

Making the appropriate substitution for $T[1]$ from (S7) results in

$$(1+r_h)T[2] = \tau + (r_s+r_h)\left((1+r_h)\tau + (r_s-r_h)T[2] + r_h(L-3)\frac{\beta}{\alpha} + 2r_hC_1\psi\right)$$
$$+ (r_s-r_h)T[3] + r_h(L-3)\frac{\beta}{\alpha} + 2r_hC_1\psi$$

This equation can be rearranged into the form

$$\left(1+r_h-r_s^2+r_h^2\right)T[2] = \tau\left(1+(r_s+r_h)(1+r_h)\right)+(1+r_s+r_h)r_h\left((l-3)\frac{\beta}{\alpha}+2C_1\psi\right)+(r_s-r_h)T[3]$$

We will substitute the values of $T[2]$ and $T[3]$ using (S12) and bringing the $C_1$ terms to the left hand side gives

$$C_1\left(\left[1+r_h-r_s^2+r_h^2\right]\left(e^{2\sqrt{\alpha}}+e^{(L-2)\sqrt{\alpha}}\right)-2\psi r_h(1+r_s+r_h)-(r_s-r_h)\left(e^{3\sqrt{\alpha}}+e^{(L-3)\sqrt{\alpha}}\right)\right)=$$
$$\tau\left(1+(r_s+r_h)(1+r_h)\right)+\frac{\beta}{\alpha}\left((1+r_s+r_h)r_h(L-3)+r_s^2+r_s-(r_h+1)^2\right)$$

which implies that

$$C_1 = \frac{\tau\left(1+(r_s+r_h)(1+r_h)\right)+\frac{\beta}{\alpha}\left((1+r_s+r_h)r_h(L-3)+r_s^2+r_s-(r_h+1)^2\right)}{\left[1+r_h-r_s^2+r_h^2\right]\left(e^{2\sqrt{\alpha}}+e^{(L-2)\sqrt{\alpha}}\right)-2\psi r_h(1+r_s+r_h)-(r_s-r_h)\left(e^{3\sqrt{\alpha}}+e^{(L-3)\sqrt{\alpha}}\right)}$$

For ease in further calculations let us assume

$$\phi_1 = 1+(r_s+r_h)(1+r_h), \qquad \phi_2 = (1+r_s+r_h)r_h(L-3)+r_s^2+r_s-(r_h+1)^2$$

$$\phi_3 = \left[1+r_h-r_s^2+r_h^2\right]\left(e^{2\sqrt{\alpha}}+e^{(L-2)\sqrt{\alpha}}\right)-2\psi r_h(1+r_s+r_h)-(r_s-r_h)\left(e^{3\sqrt{\alpha}}+e^{(L-3)\sqrt{\alpha}}\right)$$

and rewrite $C_1$ as

$$C_1 = \frac{\phi_1\tau+\frac{\phi_2\beta}{\alpha}}{\phi_3} \tag{S8}$$

Now we can evaluate for $\beta = \frac{\tau + r_h\sum_{x=1}^{L-1}T[x]}{r_s-r_h}$ as follows

$$(r_s-r_h)\beta = \tau + r_h\sum_{x=1}^{L-1}T[x]$$
$$(r_s-r_h)\beta = \tau + r_h\sum_{x=2}^{L-2}T[x] + r_h(T[1]+T[L-1])$$
$$(r_s-r_h)\beta = \tau + r_h\left((L-3)\frac{\beta}{\alpha}+2C_1\psi\right)+2r_hT[1]$$

substitute values of $C_1$, $T[1]$ and $T[2]$ to get

$$(r_s-r_h)\beta = \tau + r_h\left((L-3)\frac{\beta}{\alpha}+2\left[\frac{\phi_1\tau+\frac{\phi_2\beta}{\alpha}}{\phi_3}\right]\psi\right)$$
$$+2r_h((1+r_h)\tau+(r_s-r_h)\left(\frac{\beta}{\alpha}+\left[\frac{\phi_1\tau+\frac{\phi_2\beta}{\alpha}}{\phi_3}\right](e^{2\sqrt{\alpha}}+e^{(L-2)\sqrt{\alpha}})\right)+r_h(L-3)\frac{\beta}{\alpha}+2r_h\left[\frac{\phi_1\tau+\frac{\phi_2\beta}{\alpha}}{\phi_3}\right]\psi)$$

Taking $\beta$ terms to LHS gives

$$\beta\left[r_s - r_h - \frac{r_h(L-3)(1+2r_h)}{\alpha} - \frac{2r_h\phi_2\psi(1+2r_h)}{\alpha\phi_3} - \frac{2r_h(r_s-r_h)}{\alpha}\left(1+\frac{\phi_2(e^{2\sqrt{\alpha}}+e^{(L-2)\sqrt{\alpha}})}{\phi_3}\right)\right] =$$
$$\tau\left(1+2r_h(1+r_h)+\frac{2r_h\phi_1\psi(1+2r_h)}{\phi_3}+2r_h(r_s-r_h)\frac{\phi_1(e^{2\sqrt{\alpha}}+e^{(L-2)\sqrt{\alpha}})}{\phi_3}\right)$$

Which implies $\beta$ is given by the equation

$$\beta = \frac{\tau\left(1+2r_h(1+r_h)+\frac{2r_h\phi_1\psi(1+2r_h)}{\phi_3}+2r_h(r_s-r_h)\frac{\phi_1(e^{2\sqrt{\alpha}}+e^{(L-2)\sqrt{\alpha}})}{\phi_3}\right)}{r_s-r_h-\frac{r_h(L-3)(1+2r_h)}{\alpha}-\frac{2r_h\phi_2\psi(1+2r_h)}{\alpha\phi_3}-\frac{2r_h(r_s-r_h)}{\alpha}\left(1+\frac{\phi_2(e^{2\sqrt{\alpha}}+e^{(L-2)\sqrt{\alpha}})}{\phi_3}\right)} \tag{S9}$$

With all the unknowns identified in terms of known variables, we can write the closed-form solution in the domain $p_u \in (0,1)$ as

$$T[x] = \frac{\beta}{\alpha} + \left(\frac{\phi_1\tau + \frac{\phi_2\beta}{\alpha}}{\phi_3}\right)(e^{x\sqrt{\alpha}} + e^{(L-x)\sqrt{\alpha}}) \tag{S10}$$

This solution is in the domain $x \in [2, L-2]$ but we can calculate values at $T[1]$ and $T[L-1]$ by using (S7).

##### S3.2.1 Small $p_u$ approximation

The general solution of our differential equation is given by

$$T[x] = \frac{\beta}{\alpha} + C_1e^{x\sqrt{\alpha}} + C_2e^{-x\sqrt{\alpha}} \tag{S11}$$

For $p_u << 1/L$ we can make the assumption that $r'_s = r_s$ and $r'_h = r_h$, this would help us remove the discontinuity observed at positions $i \in \{1, L-1\}$
**Boundary condition 1:** Same as in the previous case the first derivative of $T$ at $x = \frac{L}{2}$ is zero

$$\Rightarrow \frac{\partial T[L/2]}{\partial x} = \sqrt{\alpha}\left(C_1e^{\frac{\sqrt{\alpha}L}{2}} - C_2e^{-\frac{\sqrt{\alpha}L}{2}}\right) = 0$$

from the above equation we can derive $\left[C_2 = C_1e^{\sqrt{\alpha}L}\right]$ and substitute in (S11) to get

$$T[x] = \frac{\beta}{\alpha} + C_1(e^{x\sqrt{\alpha}} + e^{(L-x)\sqrt{\alpha}}) \tag{S12}$$

**Boundary condition 2:** We have defined $T[0] = 0$, using this condition

$$T[0] = \frac{\beta}{\alpha} + C_1(1 + e^{L\sqrt{\alpha}}) = 0$$
$$\Rightarrow C_1 = \frac{-\beta}{\alpha(1+e^{L\sqrt{\alpha}})}$$

Now the only remaining unknown is $\beta$ we will evaluate this making use of the information

$$\beta = \frac{\tau + r_h\sum_{x=1}^{L-1}T[x]}{r_s - r_h} \tag{S13}$$

The summation term can be evaluated as follows

$$\sum_{x=1}^{L-1} T[x] = \sum_{x=1}^{L-1}\left[\frac{\beta}{\alpha} + \frac{-\beta}{\alpha(1+e^{L\sqrt{\alpha}})}(e^{x\sqrt{\alpha}} + e^{(L-x)\sqrt{\alpha}})\right]$$

$$= \frac{\beta}{\alpha}[L - 1 - \frac{1}{1+e^{L\sqrt{\alpha}}}(e^{\sqrt{\alpha}} + e^{2\sqrt{\alpha}} + .... + e^{(L-2)\sqrt{\alpha}}$$

$$+ e^{(L-1)\sqrt{\alpha}} + e^{(L-1)\sqrt{\alpha}} + e^{(L-2)\sqrt{\alpha}} + .... + e^{2\sqrt{\alpha}} + e^{\sqrt{\alpha}})]$$

$$= \frac{\beta}{\alpha}\left[L - 1 - \frac{2(e^{\sqrt{\alpha}} + e^{2\sqrt{\alpha}} + .... + e^{(L-2)\sqrt{\alpha}} + e^{(L-1)\sqrt{\alpha}})}{1+e^{L\sqrt{\alpha}}}\right]$$

Using geometric progression series summation

$$\sum_{x=1}^{L-1} T[x] = \frac{\beta}{\alpha}\left[L - 1 - \frac{2e^{\sqrt{\alpha}}(e^{\sqrt{\alpha}(L-1)} - 1)}{(1+e^{L\sqrt{\alpha}})(e^{\sqrt{\alpha}} - 1)}\right]$$

We will denote

$$\rho = L - 1 - \frac{2e^{\sqrt{\alpha}}(e^{\sqrt{\alpha}(L-1)} - 1)}{(1+e^{L\sqrt{\alpha}})(e^{\sqrt{\alpha}} - 1)} \tag{S14}$$

and rewrite the equation as

$$\sum_{x=1}^{L-1} T[x] = \frac{\beta}{\alpha}\rho \tag{S15}$$

Substituting the summation term in the equation for $\beta$

$$\beta = \frac{\tau + r_h\frac{\beta}{\alpha}\rho}{r_s - r_h}$$

$$\Rightarrow \beta = \frac{\tau}{r_s - r_h\left(1 + \frac{\rho}{\alpha}\right)}$$

Hence, we have shown that when $p_u$ takes values which are much less than $\frac{1}{L}$

$$T[x] = \frac{\tau}{\alpha\left(r_s - r_h\left(1 + \frac{\rho}{\alpha}\right)\right)}\left[1 - \frac{e^{x\sqrt{\alpha}} + e^{(L-x)\sqrt{\alpha}}}{1+e^{L\sqrt{\alpha}}}\right]$$

$$= \frac{\tau}{\alpha(r_s - r_h) - r_h\rho}\left[1 - \frac{e^{x\sqrt{\alpha}} + e^{(L-x)\sqrt{\alpha}}}{1+e^{L\sqrt{\alpha}}}\right]$$

where $\rho$ is given by (S14).

##### S3.2.2 Large $p_u$ approximation

For larger $p_u$ values we can make a simple approximation that $r_s \approx r_h$ and $T[2] = T[3] = ... = T[L-2]$. Applying these to the differential (S2) gives

$$-(1 + 3r_h - 2r_s)\,T[x] + (\tau + r_h(L-3)T[x] + 2r_h(T[1])) = 0 \tag{S16}$$

But from the master equation

$$T[1] = \tau + r_s'T[2] + r_h'(T[3] + T[4] + ... + T[L-1]) \tag{S17}$$

Using the same approximations $r_s \approx r_h$ and $T[2] = T[3] = ... = T[L-2]$ we can evaluate the above expression to get

$$T[1] = \frac{\tau + (r'_s + (L-4)r'_h)T[x]}{1 - r'_h} \tag{S18}$$

Substituting in (S16) and solving for T[x] yields

$$T[x] = \frac{(1 + \frac{2r_h}{1-r'_h})\tau}{1 - 2r_s - r_h(L-6) - \frac{2r_h(r'_s+(L-4)r'_h)}{1-r'_h}} \tag{S19}$$

Assuming $\tau = 1$ the above equation can be simplified by using the definition of $r_s, r'_s, r_h$ and $r'_h$ in terms of $L$ and $p_u$ be rewritten as

$$T[x] = 2 + (L-3)p_u + \frac{(2 + (L-4)p_u)^2}{2p_u} \tag{S20}$$

##### S3.2.3 The effective medium approach at large $p_u$

The Erdős–Rényi (ER) network is a mathematical model of random graphs in graph theory. In this model, each pair of nodes in a graph is connected with a certain probability, and the resulting graph is a random realization of this probability. In the high-$p_u$ limit, we take the nodes $i \in [1, L-1]$ of polymer domain to be an Erdős–Rényi network with $L-1$ nodes and calculate mean time to reach boundary nodes, closely following the approach described in [19].

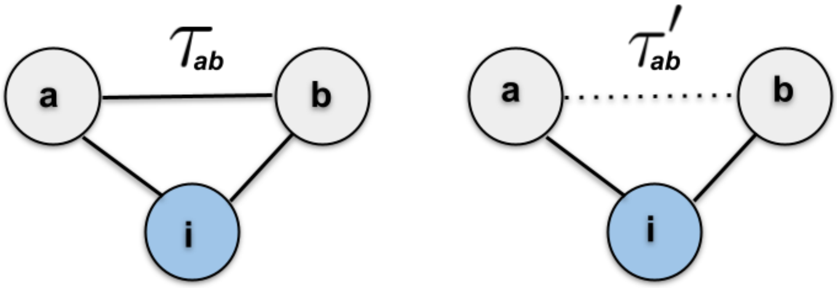


Figure S5: **Schematic of Effective medium approach:** We utilize an Erdős–Rényi network model to represent a highly compact polymer domain. The mean transit time for a random walker to travel from point a to point b is denoted as $\tau_{ab}$ when there exists a direct path connecting a and b (left). In cases where no direct path links a and b, the random walker can reach b through an intermediary point i (right). The corresponding mean transit time is then denoted as $\tau'_{ab}$.

Let $\tau_{ab}$ be the mean transit time to reach any arbitrary node $b$ starting from $a$ by a random walker in this Erdős–Rényi network with $L-1$ nodes having connection probability $p_u$ under the case where $a$ and $b$ is directly connected by a path. The number of paths coming out of $a$ excluding the adjacent nodes are $(L-4)p_u$, the adjacent nodes are always connected. The probability to reach $b$ starting from $a$ in a single step using the direct connection is $\frac{1}{2+(L-4)p_u}$. On the other hand, if the walker avoids the direct path between $a$ and $b$, the time is given by $\left(1 - \frac{1}{2+(L-4)p_u}\right)\tau'_{ab}$. Here $\tau'_{ab}$ is the average time to reach $b$ from $a$ in absence of direct path. Then we have [19],

$$\tau'_{ab} = p_u(\tau_{ab} + 1) + (1 - p_u)(\tau'_{ab} + 1) \tag{S21}$$

This allows to write a recursion relation of $\tau_{ab}$–

$$\tau_{ab} = \frac{1}{2+(L-4)p_u} + \left(1 - \frac{1}{2+(L-4)p_u}\right)\left[p_u(\tau_{ab}+1) + (1-p_u)(\tau'_{ab}+1)\right] \tag{S22}$$

Solving these two equations we get–

$$\tau_{ab} = L - 3 + \frac{1}{p_u} \qquad\qquad \tau'_{ab} = L - 3 + \frac{2}{p_u} \tag{S23}$$

Hence, the average time taken by the walker to reach $b$ starting from $a$ in a Erdős–Rényi network is–

$$\langle T_{ab}\rangle_{ER} = p_u\tau_{ab} + (1-p_u)\tau'_{ab} = L - 4 + \frac{2}{p_u} \tag{S24}$$

We are only concerned about the time it takes to reach either one of our near boundary nodes, that is 1 or $L$, assuming b to be either one of these we can argue that

$$\langle T\rangle_{ER} = \frac{(L-4)p_u + 2}{2p_u} \tag{S25}$$

We define $\langle T\rangle$ to be the mean time taken to make an exit from the chain. In this case, $\langle T\rangle$ follows the following relation–

$$\langle T\rangle = \frac{1}{2+(L-3)p_u} + \left(1 - \frac{1}{2+(L-3)p_u}\right)\left[1 + \langle T\rangle_{ER} + \langle T\rangle\right] \tag{S26}$$

Substituting $\langle T\rangle_{ER}$ and solving for $\langle T\rangle$ gives us

$$\langle T\rangle = 1 + \frac{(1+(L-3)p_u)(2+(L-2)p_u)}{2p_u} \tag{S27}$$

This estimate of mean search time matches well with the simulation for the higher $p_u$ ($p_u \to 1$) (Fig. S6). However for smaller $p_u$ values, the effective medium approach breaks down due to decreased connectivity of the network.

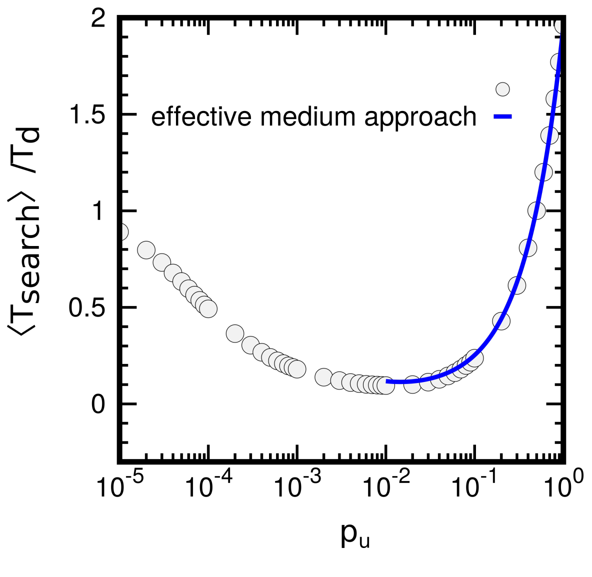


Figure S6: Scaled mean search time ($\frac{\langle T_{\text{search}}\rangle}{T_d}$) in uniformly connected network domain vs probability of contact ($p_u$) between any two non-neighbouring beads for $L = 100$. Simulation (circle) and effective medium approach Eq. S27 (blue line).

##### S3.3 The effect of 3D unbinding on mean search time

The general facilitated diffusion model proposes bulk diffusion bulk diffusion as one of the major modes of protein search. However, experiments have shown that the percentage of the time that the protein spends bound to DNA is greater than 70% (See table S1). This is very different from the most optimal conditions predicted by the facilitated diffusion model.

| **Protein name** | **Percentage of time spent bound to DNA** | **Reference** |
|---|---|---|
| RNA Polymerase | 93 | [20] |
| Lac repressor | 96 | [20] |
| Lac repressor | 87 | [6] |
| RNA Polymerase | 87.2 | [2] |
| P53 | 72 | [10] |
| P53 | 99.9 | [21] |

Table S1: Experimental quantification of 1D motion of protein during search processes

General facilitated diffusion models do not consider the polymer organisation of chromatin. In order to compare the effects of both 3D diffusion and polymer compaction in regulating search times, we calculate the mean search time in the presence of both of these effects. For dynamic polymer configurations, the mean search time can be described the recursion relation,

$$\mathbb{T}_i = \tau + \frac{1-p_{\text{off}}}{N_i}\left[\mathbb{T}_{i-1} + \mathbb{T}_{i+1} + \sum_{j\neq nn,0,L} c|i-j|^{-\gamma}\mathbb{T}_j\right] + p_{\text{off}}\mathbb{T}_f \tag{S28}$$

When the walker is at position $i$ the dynamic average of possible connections

$$N_i = 2 + \sum_{j\neq nn,0,L} c|i-j|^{-\gamma} \quad \forall\, i,j \in [1, L-1]$$

and $\mathbb{T}_f$ is the search time with polymer starting in the bulk and is defined as,

$$\mathbb{T}_f = \tau_f + \frac{\sum_{j=0}^{L}\mathbb{T}_j}{L+1}$$

By introducing $p_{\text{off}} \neq 0$, we allow protein to go to solution with unbinding probability $p_{\text{off}}$. Upon unbinding, the protein rebinds randomly to any bead on chromatin after a characteristic time $\tau_f$. We assume $\tau_f = 100\tau$ [23], although our results do not depend on the specific choice of $\tau_f$. When unbinding probability is low, the polymer compaction can lower the search times by allowing intersegmental transfer. There exists an optimal compaction for which search time is minimum. However for high $p_{\text{off}}$, the polymer structure plays no role, resulting constant search times for all level of compaction (Fig. S7a,c).

On the other hand, when the polymer compaction is low (low $p_u$ or, high $\gamma$), there is optimal unbinding probability for which search time is minimum. However, the non-monotonic behaviour is lost for higher compaction level, which is more relevant for TAD like chromatin domains (Fig. S7b,d).

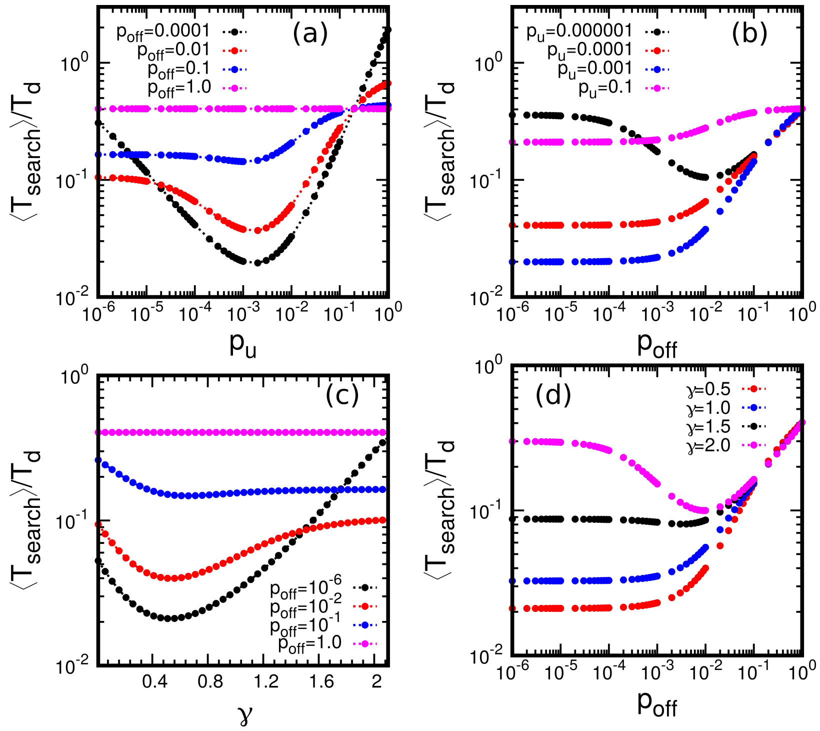


Figure S7: Average search time as function of polymer compaction for varying $p_{\text{off}}$ in **(a)** uniformly connected domain, **(c)** power-law connected domain. Average search time as function of unbinding probability $p_{\text{off}}$ for varying compaction in **(b)** uniformly connected domain, **(d)** power-law connected domain.

#### S4 Search process on FRC, LJ and soft LJ polymer

We investigate the persistence of the non-monotonic behavior of $\langle T_{\text{search}}\rangle$ when canonical polymer models are used to represent chromatin structure of a TAD. To begin, we construct polymer structures using different models. These structures are then converted into a network representation ($c_{ij}$). This conversion process involves defining a cut-off radius ($r_c$). If the 3D distance ($r_{ij}$) between two non-bonded beads $(i, j)$ in the polymer structure is within the cut-off radius, we consider them to be in contact ($c_{ij} = 1$).

$$c_{ij} = \begin{cases} 1 & r_{ij} \leq r_c \\ 0 & r_{ij} > r_c \end{cases} \tag{S29}$$

The connectivity along chain is ensured by employing the condition $c_{i,i+1} = 1$. To characterise the compactness of the polymer we calculate radius of gyration ($R_g$) of the polymer, which is defined as-

$$R_g = \sqrt{\frac{1}{N}\sum_{i=1}^{N}(\mathbf{r}_i - \mathbf{r}_{\text{CM}})^2}$$

where $N$ is the number of monomers in the polymer chain, $r_i$ is the position vector of the i-th monomer, $r_{CM}$ is the position vector of the center of mass of the polymer, given by $r_{\text{CM}} = \frac{1}{N}\sum_{i=1}^{N} r_i$. In addition, we also calculate total number of non-neighbouring bonds ($N_b$), which is defined as-

$$N_b = \sum_{i>j+1}^{N} c_{ij}$$

Finally, we perform random walks on these obtained network structures to explore the dynamics of protein and analyze the behavior of $\langle T_{\text{search}} \rangle$ in relation to the polymer compactness.

##### S4.1 Freely rotating chain model

We first study the Freely Rotating Chain (FRC) model which has been used to investigate chromatin behaviour [17, 1]. Neighbouring bonds in an FRC polymer are constrained to lie on a cone subtending an angle $\theta$ [16]. The angle $\theta$ controls the compaction of the polymer, with large $\theta$ leading to a collapsed state (Fig. S8a). As the polymer collapses, the number of non-neighbouring bonds $N_b$ increases monotonically ($r_c = 1.2\sigma$) (Fig. S8b), indicating a greater probability of intersegmental jumps. We simulated search on FRC networks, and the mean search times $\langle T_{\text{search}} \rangle$ shows a non-monotonic behaviour with increasing bond angle (see main text). This can be also interpreted as a function of polymer compactness or collapse. Thus in line with the expectations from the network model of polymer, there exists an optimal compaction where search times are minimum.

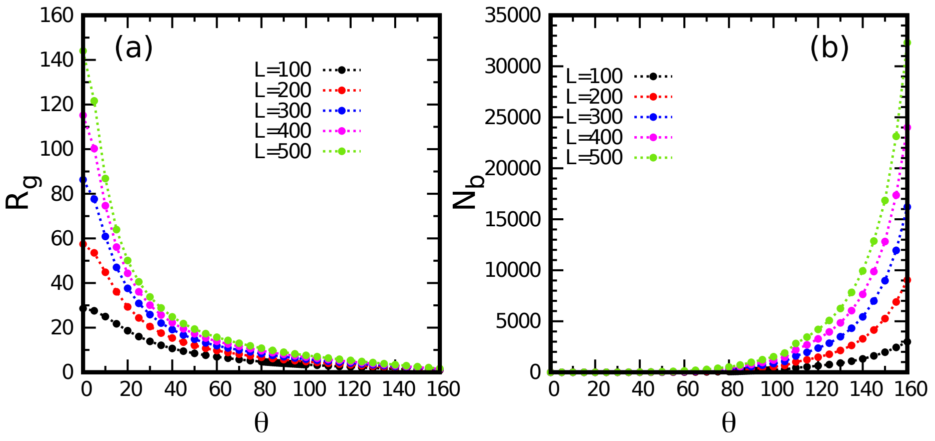


Figure S8: **a.** Radius of gyration ($R_g$) as a function of FRC chain angle ($\theta$). **b.** Number of inter-segmental contacts ($N_b$) as a function of FRC chain angle ($\theta$).

##### S4.2 LJ bead-spring polymer model

We next use a bead-spring polymer model of chromatin with attractive Lennard-Jones (LJ) potential [3, 4, 18]. This introduces self-avoidance effects, and bond length fluctuations absent in the FRC model. Any pair of beads of diameter $\sigma$, separated by distance $r$, interacts via the potential,

$$E_{LJ}(r) = \begin{cases} 4\epsilon \left[ \left(\frac{\sigma}{r}\right)^{12} - \left(\frac{\sigma}{r}\right)^{6} \right] & r < 2.5\sigma \\ 0 & r \geq 2.5\sigma \end{cases} \tag{S30}$$

###### Simulation of Lennard-Jones polymer

To model chromatin we have used a bead-spring polymer consisting of $L+1$ beads each of size (diameter) $\sigma$. The total energy ($E$) of the polymer is given by-

$$E = \sum_{i=1}^{L-1} E_{spring}(|\vec{r}_{i+1} - \vec{r}_i|) + \sum_{i=1}^{L-1} \sum_{j=i+1}^{L} E_{LJ}(|\vec{r}_j - \vec{r}_i|) \tag{S31}$$

Here $\vec{r}_i$ is the position vector of $i^{th}$ bead. The first term in Eqn. S31 denotes energy contribution ($E_{spring}(r) = \frac{k}{2}(r-\sigma)^2$) from the harmonic springs which connect adjacent beads of the polymer along the chain. The second term in Eqn. S31 represents the attractive interactions via Lennard-Jones potential.

$$E_{LJ}(r) = \begin{cases} 4\epsilon\left[\left(\frac{\sigma}{r}\right)^{12} - \left(\frac{\sigma}{r}\right)^{6}\right] & r < 2.5\sigma \\ 0 & r \geq 2.5\sigma \end{cases} \tag{S32}$$

Here, $\epsilon$ is the strength of the attractive interaction. The simulations were performed using reduced units where we set mass of the bead $m = 1$, diameter of the bead $\sigma = 1$ and energy is measured in units of $k_B T$. The value of spring constant is $k = 60$. We varied the LJ interaction strength $\epsilon$ to simulate polymers of different compaction. We chose the time step of $\Delta t = 0.01$ and damp parameter in LAMMPS [22] was taken as 1. We performed molecular dynamics simulations with an implicit solvent using a scheme known as Langevin dynamics which follows the Eq. S33, where the last term of represents the random collisions caused by the solvent particles.

$$m\frac{d^2\mathbf{r}}{dt^2} = -\nabla E(\mathbf{r}) - \gamma\frac{d\mathbf{r}}{dt} + \sqrt{2\gamma k_B T}\boldsymbol{\eta}(t) \tag{S33}$$

We constructed a 1000 distinct bead-spring polymer configurations, each having $L + 1$ beads.

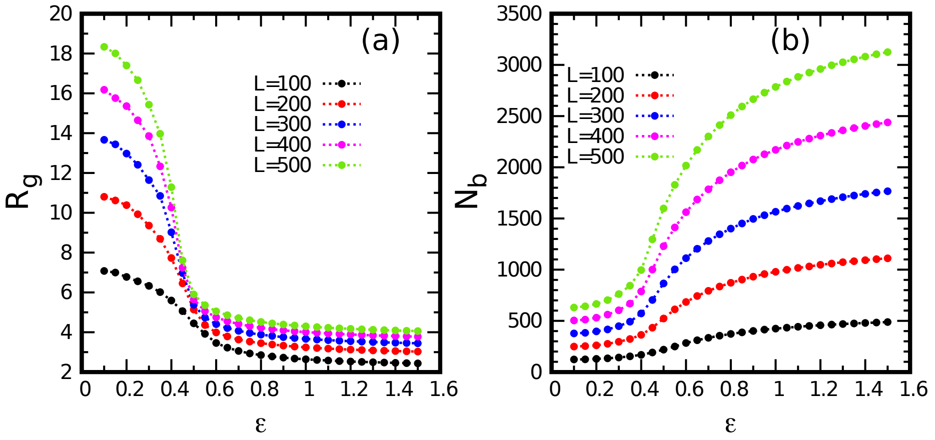


Figure S9: **a.** Radius of gyration ($R_g$) as a function of LJ interaction strength ($\epsilon$). **b.** Number of inter-segmental contacts ($N_b$) as a function of LJ interaction strength ($\epsilon$).

A LJ polymer undergoes a second-order phase transition from a open to a compact state (see Fig. S9a), accompanied by an increase in the number of intersegmental bonds (see Fig. S9b), as $\epsilon$ is increased ($r_c = 1.8\sigma$). The search times $\langle T_{\text{search}}\rangle$ on these LJ polymer backbones exhibited a non-monotonic pattern as the LJ interaction strength is increased. There was a distinct minimum observed in the search times (see main text Fig. 4), occurring close to the collapse transition. Thus a LJ polymer qualitatively mirrors the results obtained from the network models and FRC polymer model.

However, unlike the significant rise seen in the FRC and the network model, the extent of $\langle T_{\text{search}}\rangle$ increase beyond a critical $\epsilon$ is less pronounced in this case. This limited increase is due to the inherent hard-core repulsion in the LJ potential, which imposes a lower limit on polymer compaction, and on the number of intersegmental bonds. This interplay between LJ interactions and polymer structure thus introduces a unique feature that distinguishes it from both FRC and the network model.

##### S4.3 Soft LJ bead-spring polymer model

Is this weak non-monotonic rise in $\langle T_{\text{search}}\rangle$ then the correct physical description of search times? The answer hinges on whether the hardcore LJ potential best represents coarse-grained chromatin conformations. Experimental and theoretical studies of chromatin configurations and 3D distances suggest that a soft inter-bead potential that allows for overlap is a more realistic description of coarse-grained chromatin [9, 7].

The approach of modeling chromatin as a bead-spring polymer at a coarse-grained level may face challenges, as highlighted by previous research [9]. Specifically, there tends to be a notable degree of overlap among the constituent beads in this coarse-grained representation [9]. In response to this issue, the model deviates from employing a Lennard-Jones potential with a hardcore repulsion term. Instead, it adopts a distinct potential that incorporates a more gently repulsive component. The total energy ($E$) governing the polymer is expressed through this modified potential as-

$$E = \sum_{i=1}^{L-1} E_{spring}(|\vec{r}_{i+1} - \vec{r}_i|) + \sum_{i=1}^{L-1} \sum_{j=i+1}^{L} E_{soft}(|\vec{r}_j - \vec{r}_i|) \tag{S34}$$

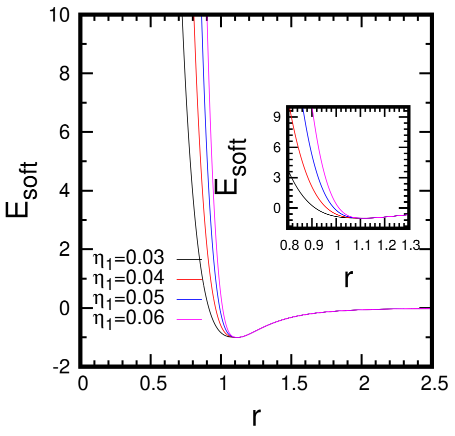


Figure S10: Nature of the soft LJ potential (eq. S35) for $\eta_1 = 0.03, 0.04, 0.05, 0.06$. $\eta_1$ controls the softness of the polymer. As $\eta_1$ is decreased softness of the polymer increases. In the inset, zoomed view of the potential near minima.

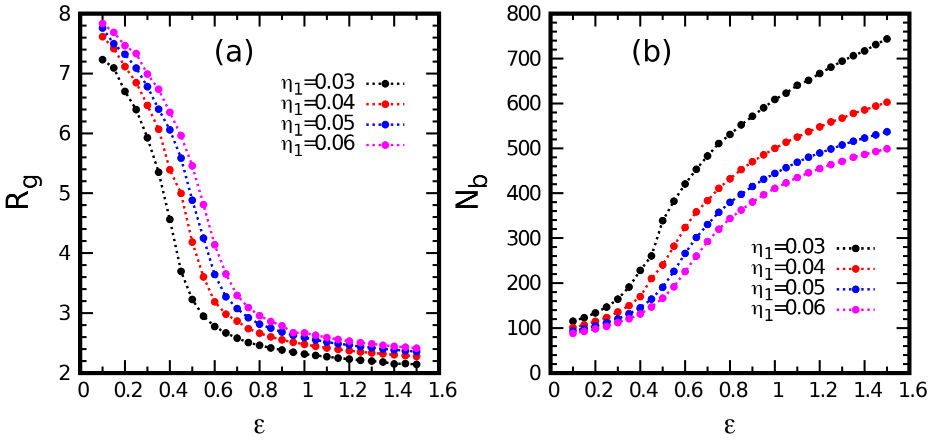


Figure S11: **a.** Radius of gyration ($R_g$) as a function of soft LJ interaction strength ($\epsilon$) for $\eta_1 = 0.03, 0.04, 0.05, 0.06$ for $L = 100$. **b.** Number of inter-segmental contacts ($N_b$) as a function of soft LJ interaction strength ($\epsilon$).

This energy eq. S34 takes into account both the bonded interactions between adjacent beads and the softer repulsive forces that aim to mitigate bead overlap. $E_{soft}$ is given by-

$$E_{soft}(r) = \begin{cases} V_0[1-(\frac{r}{r_m})^{\eta_1}]^{\eta_2} - \epsilon & 0 \leq r < r_m \\ 4\epsilon\left[\left(\frac{\sigma}{r}\right)^{12} - \left(\frac{\sigma}{r}\right)^{6}\right] & r_m \leq r < 2.5\sigma \\ 0 & r \geq 2.5\sigma \end{cases} \quad \text{(S35)}$$

The expression $V_0 - \epsilon$ denotes the energy penalty associated with complete overlap, and the parameters $\eta_1$ and $\eta_2$ are utilized to adjust the softness of the potential, as illustrated in Figure S10. When $r < r_m$, the potential becomes repulsive but with a softer behavior compared to the LJ potential. For $r_m \leq r < 2.5\sigma$, the potential aligns with the LJ potential, inducing attractive interactions. Beyond a cutoff distance ($r \geq 2.5\sigma$), the interaction energy is set to zero. Parameters of the potential for our study: $V_0 = 10^7$, $\eta_1 =$ variable, $\eta_2 = 3.16$, $r_m = 1.12$, $\sigma = 1$.

where $V_0 - \epsilon$ is the energy penalty at complete overlap, and $\eta_1$ and $\eta_2$ tune the softness of the potential. We investigate $\langle T_{\text{search}}\rangle$ in polymers with varying degrees of softness by tuning $\eta_1$. A lower value of $\eta_1$ indicates a softer polymer, whereas high $\eta_1$ resembles the classical LJ potential. For a very soft polymer ($\eta_1 = 0.03$), $\langle T_{\text{search}}\rangle$ shows a strong non-monotonicity, analogous to the FRC and the network results. This is because a soft polymer core cannot effectively enforce volume exclusion, resulting in a more tightly packed polymer compared to the conventional LJ model (Fig. S11a). This then leads to a higher number of non-neighboring contacts ($r_c = 1.8\sigma$) within the polymer domain (Fig. S11b). As we decrease the softness (increasing $\eta_1$), the degree of non-monotonicity in the search times decreases and approaches the LJ-like behaviour. Thus non-monotonic behaviour of search times remains a crucial feature of protein diffusion on coarse-grained chromatin-like soft polymer topologies. The degree of non-monotonicity $\langle T_{\text{search}}\rangle$ are controlled by the softness of the inter-bead potential.

#### S5 Protein motion in coarse-grained chromatin

Proteins within the nucleus search for their specific target sites and bind to them. These target sites typically consist of specific sequences of a few base pairs. In computational modeling, we often coarse-grain chromatin to a lower resolution. In this coarse-grained representation, each chromatin bead represents several base pairs of DNA. When modeling the search process within coarse-grained chromatin, the description of the search strategies needs to be adjusted to account for the coarse-graining level. One key aspect of interest is the ratio of the time a protein spends bound to chromatin versus freely diffusing in the bulk nuclear environment.

While the widely studied facilitated diffusion model suggests that 1D motion along a chain accompanied by 3D bulk exploration is an efficient search strategy, bulk diffusion may not be a viable description for motion on coarse-grained chromatin at physiological chromatin volume fractions. The facilitated diffusion model is proposed for prokaryotic systems, where the density of chromatin is relatively low. In eukaryotic cells, the volume fraction of chromatin is, on average, 10 percent. However, chromatin is not uniformly distributed in the nucleus, and the local volume fraction within Topologically Associating Domains (TAD) can reach up to 35 percent [14]. The high local density of chromatin restricts the bulk exploration of proteins, and hence the protein stays non-specifically bound to DNA for long periods of time. Further, the coarse-grained description of chromatin means that a unit bead contains a few kilobase pairs (kbp) of DNA, and the bead size becomes of the order of 10-100 nanometers. Coarse-grained beads are also prone to overlapping. Therefore, the 3D diffusion at the base pair level is subsumed into the effective 1D motion in the coarse-grained picture.

In this section, we aim to determine the upper limit of the ratio of time a protein spends freely diffusing in the bulk nuclear environment versus the time it spends bound to chromatin. To achieve this, we employ a bead-spring polymer model consisting of 3000 beads within a variable-sized box, allowing us to control the volume fraction of the system. Each bead in the polymer represents a 200bp chromatin segment, which is the experimentally known smallest unit of chromatin. The protein is represented as a single particle moving randomly within the box, with all interactions between all beads (protein+polymer) set to be purely self-avoiding. We conduct simulations of this system using LAMMPS and collect trajectories of the protein. Upon obtaining the trajectories, we coarse-grain the polymer using the following strategy: we consider five consecutive beads along the polymer chain and calculate the end-to-end vector for this segment. We then place a coarse-grained bead at the center of mass of these five beads, with a diameter equal to the calculated end-to-end distance of this segment. Subsequently, we determine the distance of the protein from the center of each coarse-grained bead. If the protein is within a cutoff radius, we consider it to be bound to the chromatin. Otherwise, we consider the protein to be freely moving in the bulk nuclear environment.

In figure S12, we plot residence probability in 3D diffusion mode i.e. when the protein is not bound to chromatin as a function of the volume fraction of the chromatin in the box. As the volume fraction is increased, the bulk exploration is decreased due to lack of space for free movement in a highly dense chromatin region. This indicates, even if there is possibility that protein is following 1D+3D search strategy, that would not show up significantly while modeling the search process using a coarse-grained description. The motion of a protein can be purely modeled as a combination of slides and intersegmental jumps in such scenario. It is also important to note, in our analysis, we have not considered any specific or non-specific attractive interactions between the protein and the chromatin polymer. Additionally, we have not taken into account the structural constraints imposed by the chromatin polymer itself. In reality, both of these factors would further limit the protein's bulk exploration. For example, specific interactions between the protein and the chromatin polymer could result in the protein preferentially binding to certain regions of the chromatin, reducing its overall exploration of the bulk. Similarly, the structural constraints of the chromatin polymer, such as loops and domains, would create barriers that the protein would need to navigate around, further limiting its exploration. Overall, the plot shows an estimate of upper limit of 3D diffusion probability in compact TAD like domains.

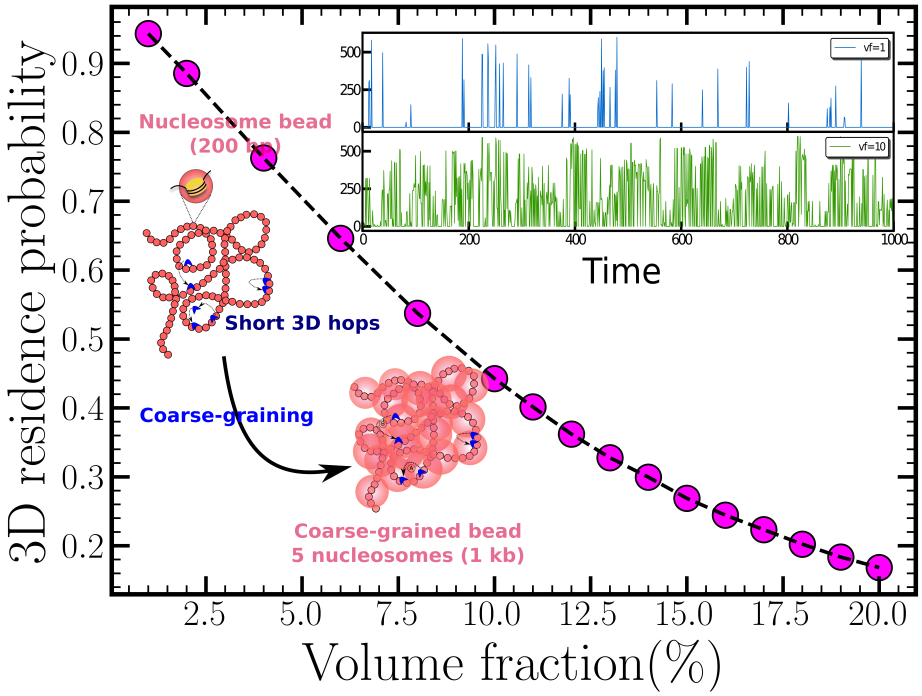


Figure S12: Residence probability in 3D diffusion mode, where the protein is not bound to chromatin, varies as a function of the volume fraction of chromatin in the box. As the volume fraction increases, bulk exploration decreases due to the limited space for free movement in densely packed chromatin regions. The inset depicts the protein's trajectory for volume fractions of 1 and 10. At low volume fractions, motion is primarily governed by binding and unbinding processes. However, at high volume fractions, the motion exhibits numerous intersegmental jumps.

#### S6 Dynamics of protein in network constructed using Hi-C data

To examine the proteins search process in real chromatin architecture we used Hi-C experiment data from [15]. The primary Hi-C map used in our analyses was a one-kilobase resolution map, generated in the GM12878 human lymphoblastoid cell line. Hi-C measures the contact frequencies between chromatin segments in 3D space, providing insights into the organization of chromatin. We scale the KR normalized [11] Hi-C contact frequency matrix of each chromosomes (Chr1-Chr22) by the largest off-diagonal element to obtain contact probabilities ($P_{ij}$). All analyses involving TADs in this map were performed using a list of 8355 domains available from [15] annotated via the Arrowhead algorithm.

We define intra-TAD contact probability $P_c(s)$ of two points separated by genomic distance $s$ as the average contact probability between of all genomic loci separated by genomic distance $s$ within a TAD. All contact probability plots are displayed in main text on log-log axes with distance s binned logarithmically. Contact probability exhibited a power law, within a range of values of $s$. We measured $\gamma$ as the slope of the best-fit line and $c$ as the $y$-intercept, when plotted on log-log axes, within a chosen range of distances. The obtained $\gamma$ values signifies the compaction of the corresponding TAD.

To model the dynamics of a transcription factor (TF) within a real chromatin structure, we use previously described slide and intersegmental jump model on a network. The network mimics the chromatin structure and contacts within a single TAD in 3D space. The network structure is established using $P_{ij}$ values obtained from the experimental Hi-C data in the

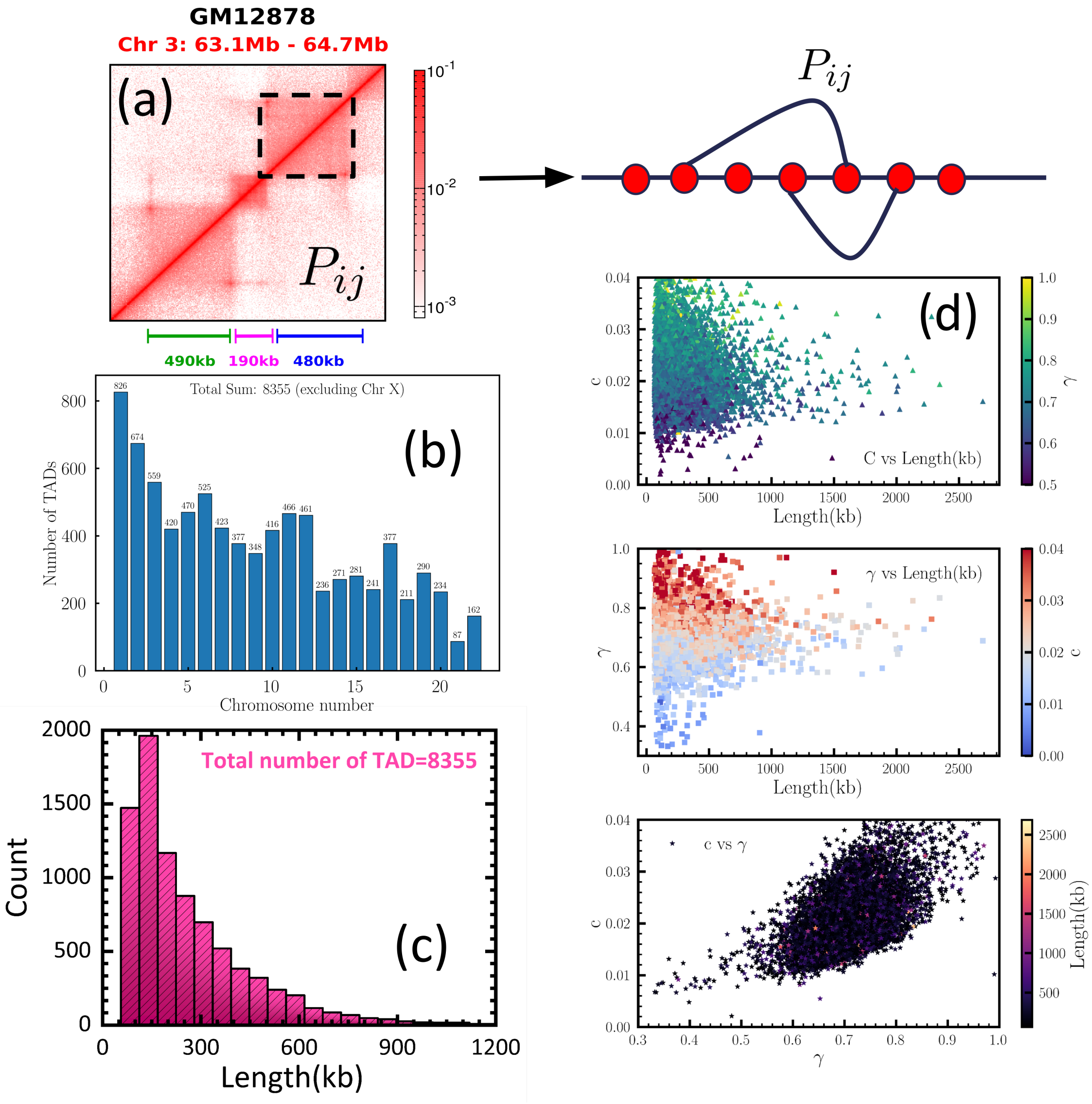


Figure S13: **(a)** Representative TAD from Human Hi-C data which provides contact probability between any two segment $i$ and $j$ as $P_{ij}$. An ensemble of networks is created using Hi-C data where contact between $i$ and $j$ with probability $P_{ij}$, extracted from Hi-C matrix. **(b)** Chromosome-wise number of TADs studied. **(c)** Distribution of lengths of TADs. **(d)** Variation of $c$, $\gamma$ and $L$ for all TADs.

following way.

$$c_{ij} = \begin{cases} 1 & r_n \leq P_{ij} \\ 0 & r_n > P_{ij} \end{cases} \tag{S36}$$

where $r_n$ is a uniform random number in [0, 1]. Here $c_{ij}$ denotes the connectivity of chromatin polymer within a TAD. Along with connectivity obtained from $P_{ij}$ using Hi-C, we also ensure $c_{i,i+1} = c_{i,i-1} = 1$ to ensure the polymer connectivity along the backbone. After constructing the network structure based on Hi-C data, we can simulate the dynamic behavior of a protein within a realistic chromatin environment. We calculate the ensemble average search times ($\langle T_{\text{search}} \rangle$) i.e. time to reach one of the TAD boundary by a random walker.
